# Alternative splicing edits intrinsically disordered regions across human protein isoforms

**DOI:** 10.64898/2026.11.23.768762

**Authors:** Tommy Phan, Susie Song, Joshua M. Akey

## Abstract

Intrinsically disordered regions (IDRs) lack a stable three-dimensional structure, yet they mediate many of the interactions and regulatory functions of proteins. Alternative splicing produces multiple protein isoforms from a single gene, and because individual exons can encode all or part of an IDR, splicing has the potential to add, remove or reshape disordered regions. However, it remains unclear whether splicing preferentially acts on disordered sequence in all kinds of proteins, how it changes an IDR from one isoform to another, and how this relationship arose during evolution. Here, we map predicted disorder onto the coding exons and genomic codons of human protein isoforms, which allows any two isoforms of a gene to be compared directly, and analyse this map across protein classes and families. We find that alternatively spliced exons are more disordered than constitutive exons in every class of protein, from transcription factors to metabolic enzymes, and that splicing usually changes IDRs at their ends, adding or removing disordered segments as modular units rather than cutting through them. The disordered sequence that splicing varies is compositionally distinct from the disorder that all isoforms share. Notably, the apparent loss of disorder over gene evolution reflects differences between protein families rather than change within them, whereas exons whose splice sites are conserved in few mammals encode more disorder than the other exons of the same gene, with a composition resembling that of young genes. Our results suggest that alternative splicing treats intrinsic disorder as a modular, evolvable layer of protein architecture, and that exons with poorly conserved splice sites may be one route by which disordered sequence enters proteins.

## 1 Introduction

We traditionally think of proteins as well-defined three-dimensional structures whose shapes dictate their functions. Yet many proteins contain intrinsically disordered regions (IDRs) that lack a sta-ble structure and instead exist as ensembles of rapidly interconverting conformations[1, 2]. Rather than being encoded in a fold, the behaviour of an IDR is encoded in its amino-acid composition and patterning[3]. This flexibility lets IDRs carry short linear motifs and modification sites that mediate transient, multivalent interactions[4], and lets them drive the formation of biomolecular condensates[5, 6]. Disorder is not evenly spread across the proteome: regulatory proteins such as transcription factors and chromatin regulators are among the most disordered, whereas enzymes and transporters are among the most ordered[7–9].

Alternative splicing adds a further layer of protein diversity. By including or skipping exons, a single gene can produce several protein isoforms, and most human multi-exon genes do so[10, 11]. These isoforms can behave as functionally distinct proteins, differing in their interaction partners and regulatory activities[12, 13]. Exons are discrete, modular units of coding sequence, and an exon can encode all or part of a disordered region. Splicing therefore has the potential to add, remove or reshape IDRs, and with them the motifs and interaction surfaces they carry. Consistent with this idea, alternatively spliced sequence is enriched for disorder across the human proteome[14–17].

Despite this, the evidence remains limited and largely dated. The foundational studies are one to two decades old, predate comprehensive isoform annotation and current disorder predictors, and examined dozens of proteins[14], tissue-regulated exons[15] or isoforms expressed in a single tissue[17] rather than every protein in the proteome. Several fundamental questions therefore remain. First, it is unclear whether the link between splicing and disorder is a general property of proteins or a feature of the highly disordered regulatory proteins that dominate proteome-wide averages. Second, it is unclear what splicing actually does to an IDR when one isoform is exchanged for another: whether disordered regions are gained and lost whole, trimmed at their ends, or interrupted in their middles. Earlier work has typically described each alternative isoform as a set of differences from one full-length protein[14], which leaves open how IDRs vary across all of a gene’s isoforms at once. Third, it is unclear how this relationship arose during evolution. Under the disorder-to-order model, new proteins begin relatively disordered and become more ordered over time[9], but it is not known whether individual proteins become more ordered as they evolve or whether old and young protein families simply differ, nor whether newly emerging exons contribute disordered sequence.

Here, we comprehensively map predicted IDRs onto the coding exons and genomic coordinates of human protein isoforms. By integrating curated proteome-wide annotations, deep-learning-based disorder prediction and exact transcript-to-genome mappings, we track every residue by the genomic codon that encodes it, which allows all isoforms of a gene to be compared globally, each against every other. Analysing this map across protein classes and families, and alongside the ages of genes and the conservation of their splice sites, we find that alternative splicing preferentially acts on disordered sequence in every kind of protein, that it remodels IDRs mostly at their ends, and that disordered sequence entering through exons with poorly conserved splice sites resembles the disorder of young genes. Together, our results suggest that splicing treats intrinsic disorder as a modular and evolvable layer of protein architecture.

## 2 Results

### 2.1 A global, exon-resolved map of disorder across human protein isoforms and classes

An overview of the question, approach and findings is in Fig. 1.

**Fig. 1.**
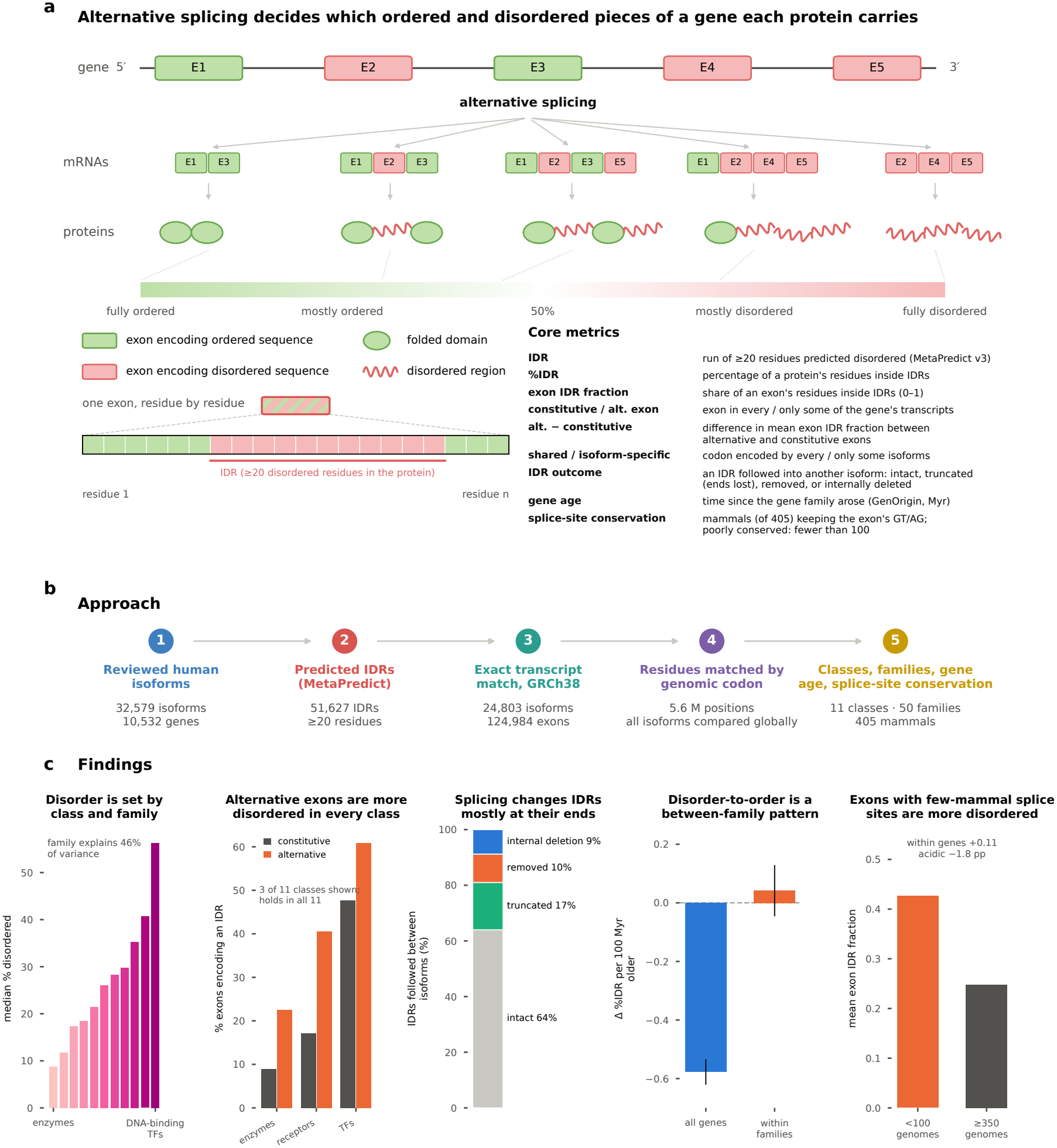
Overview: alternative splicing, intrinsic disorder and their evolution across human protein classes. **a**, A gene’s exons encode ordered (green) or disordered (pink) sequence; alternative splicing combines them into isoforms that range from fully ordered to fully disordered proteins. Bottom left, an exon that encodes both ordered and disordered sequence (striped) shown residue by residue, with the part that lies in an IDR (at least 20 consecutive residues predicted disordered in the whole protein). Bottom right, definitions of the core metrics used throughout; all terms are defined in Extended Data Table 1. **b**, Approach and the number of isoforms, IDRs, exons and positions at each step (details in Extended Data Fig. 1). **c**, Main findings: median canonical disorder across the eleven protein classes (Fig. 2c, Fig. 3); exons encoding an IDR, constitutive and alternative, in three example classes (all classes in Fig. 4); outcomes of IDRs followed between isoforms (Fig. 5); change in disorder with gene age by class and within protein families (Fig. 6a,b; Extended Data Fig. 8); mean IDR fraction of exons with poorly conserved (*<*100 of 405 mammals) and deeply conserved (*≥*350) splice sites (Fig. 7).

We started from reviewed (Swiss-Prot) human proteins with protein-level evidence and kept genes with at least two annotated isoforms: 32,579 isoforms (10,532 canonical and 22,047 alternative) from 10,532 genes, with 51,627 IDRs of at least 20 residues predicted by MetaPredict[18] (Extended Data Fig. 1; Methods). We kept isoforms whose GENCODE v50 translation is identical to the UniProt sequence and that resolve to one coding structure (24,803 isoforms) and projected every IDR and ordered segment to GRCh38 with a two-way translation check that all 76,781 segments passed. For each gene, an exon is alternative if it occurs in some but not all of the gene’s mapped transcripts and constitutive otherwise; the exon analyses use 124,984 unique coding exons from 8,196 genes.

To compare all isoforms globally, we keyed every residue of every mapped isoform by the genomic position of its codon (Fig. 2a). A position is shared if every mapped isoform of the gene encodes it and isoform-specific otherwise, and isoform-specific positions are graded by the fraction of the gene’s isoforms that include them. Two isoforms share a residue only if the same codon encodes it, so sequence from a different exon, or from the same exon read in another frame, is never counted as shared. Exon labels and codon labels mostly agree but are not the same: 99.8% of isoform-specific positions lie in alternative exons, whereas 28.5% of the positions in alternative exons are encoded by every isoform, because the exon’s boundary shifts in some isoforms (half of these positions) or because only its untranslated end, and with it the exon identifier, differs (Fig. 2b). Comparing codons therefore isolates the residues that actually differ between isoforms. This gives 5.6 million coding positions in 8,168 genes. We assigned each gene to one of eleven mutually exclusive protein classes from UniProt and HumanTFs annotations[19] (Methods), recorded its UniProt family, and dated genes with GenOrigin[20] and exons with splice-site conservation across 405 mammals[21].

**Fig. 2.**
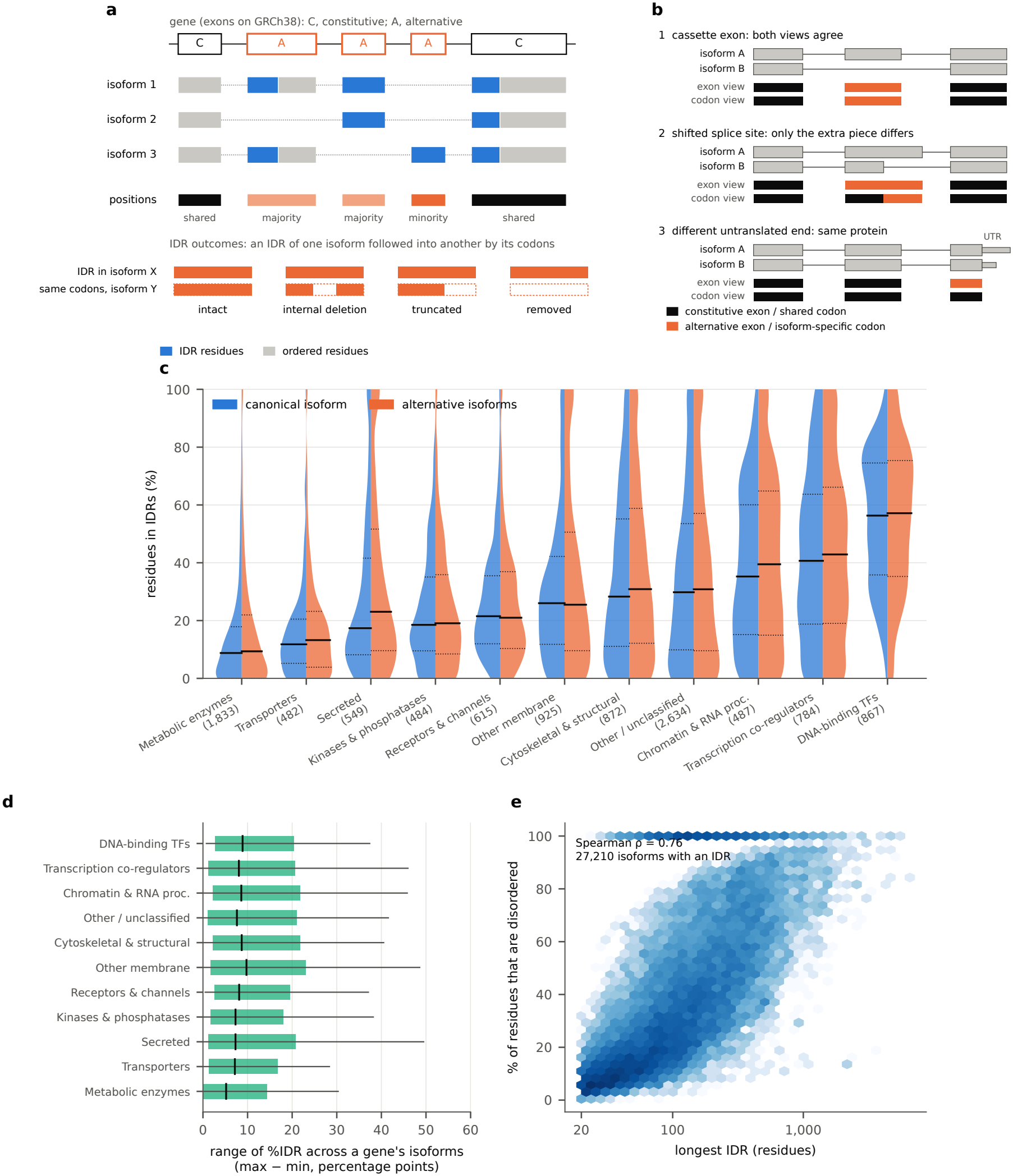
A global, exon-resolved map of disorder across human protein isoforms and protein classes. **a**, Definitions on a toy gene with three isoforms. Exons used by every mapped transcript are constitutive (C) and the others alternative (A). Residues are keyed by the genomic position of their codon: shared positions are encoded by every isoform and isoform-specific positions by some (a majority, half or a minority). Each IDR of one isoform (orange) is followed by its codons into every other isoform and classified as intact, internally deleted, truncated or removed; dashed outline, codons absent from the other isoform. **b**, Exon view versus codon view in three splicing cases. A cassette exon is alternative in the exon view and isoform-specific in the codon view. When a splice site shifts, the long and short versions of the exon are different exons, so the exon view calls all of it alternative, whereas only the extra piece is isoform-specific. When only the untranslated end of an exon differs (thin box, UTR), the exon has a new identifier and is alternative, although every codon is shared. Across 8,168 genes, 99.8% of isoform-specific positions lie in alternative exons and 28.5% of positions in alternative exons are shared. **c**, Percentage of residues in IDRs for canonical (left) and alternative (right) isoforms in eleven protein classes, ordered by median canonical disorder; number of genes in parentheses; solid lines, medians; dotted lines, quartiles. **d**, Range of disorder across each gene’s isoforms (maximum minus minimum %IDR); boxes, interquartile range; whiskers, 10th–90th percentiles. **e**, Length of the longest IDR versus disorder content for isoforms with an IDR (hexagonal bins); shading, log isoform count.

Median disorder of the canonical isoform ranged more than sixfold across classes, from 8.8% of residues in metabolic enzymes and 11.8% in transporters to 40.7% in transcription co-regulators and 56.3% in DNA-binding transcription factors (Fig. 2c). A third of metabolic enzymes (33%) had no IDR, against 3% of DNA-binding transcription factors. Within each class, canonical and alternative isoforms had similar disorder; only secreted proteins differed (median 17.4% versus 23.0%; Mann–Whitney *P* = 1.7 *×* 10^*−*3^). Isoforms of the same gene differed in disorder by a median of 7.5 percentage points, and this range was similar across classes (medians 5.2 in metabolic enzymes to 9.8 in other membrane proteins; Fig. 2d), so splicing varies disorder by a comparable amount in mostly ordered and mostly disordered proteins. Across isoforms, disorder content tracked the length of the longest IDR (Spearman *ρ* = 0.76; Fig. 2e): disordered proteins are those with long IDRs rather than many short ones.

### 2.2 Protein family and annotations resolve disorder beyond class

Protein family resolved disorder more finely than class. Among 50 UniProt families with at least 15 genes (2,076 genes), family explained 46% of the variance in canonical disorder, against 33% for class in the same genes and 2.4% for families with shuffled labels (95th percentile 3.3%; Fig. 3a). The most disordered families were the bZIP (median 77%), paired homeobox (75%) and ETS (69%) transcription factors and the SR splicing factors (66%); in the least disordered, cytochrome P450, short-chain dehydrogenase/reductase and sulfotransferase enzymes, the median protein had no IDR. Families within a class could differ widely: among DNA-binding transcription factors, the C2H2 zinc-finger family, whose proteins are dominated by arrays of folded zinc fingers, was less disordered (40%) than the bZIP, homeobox or ETS families, and the nuclear hormone receptors less still (32%).

**Fig. 3.**
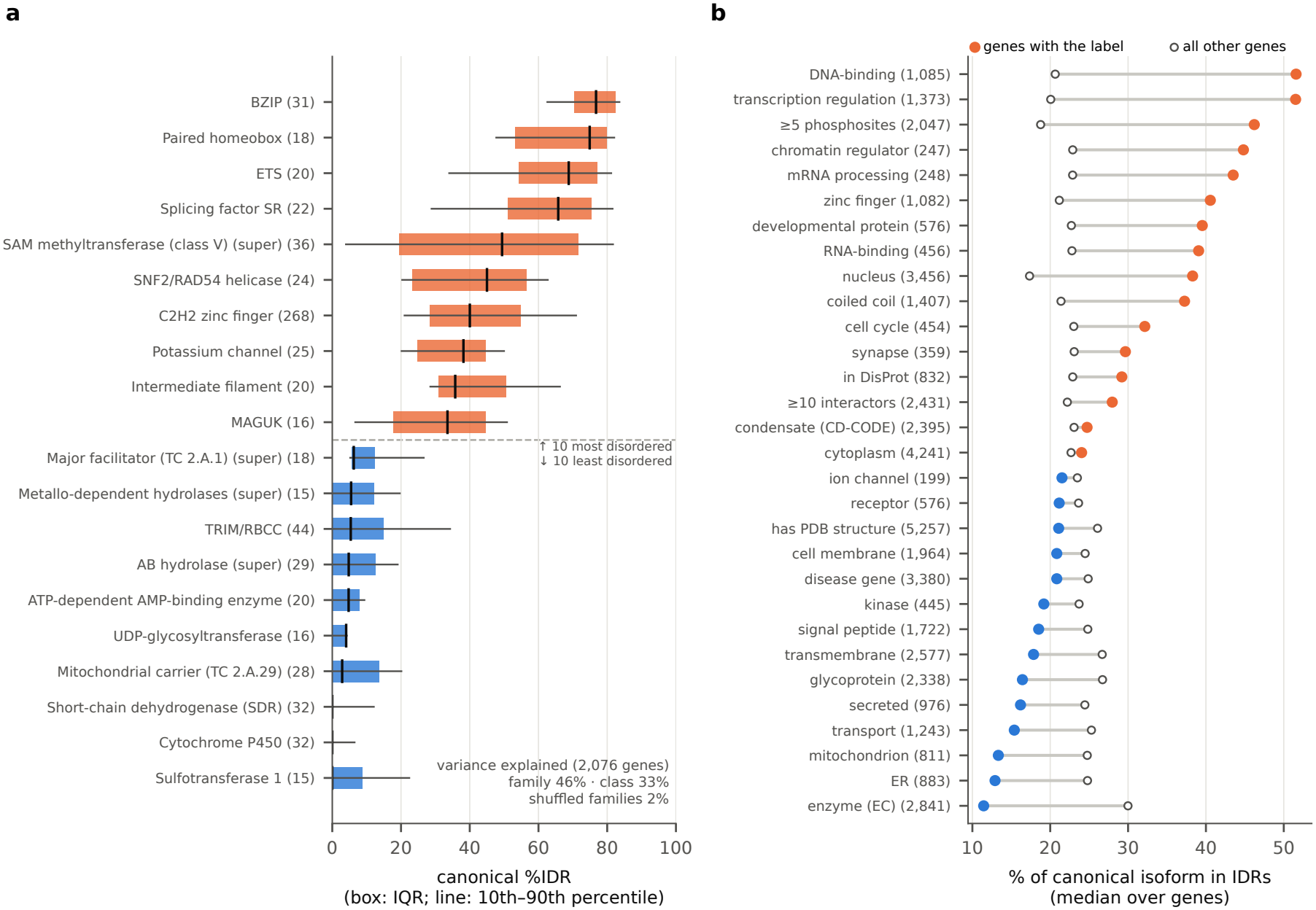
Protein families and annotations resolve disorder beyond broad class. **a**, Canonical disorder of the ten most and ten least disordered UniProt families with at least 15 genes; boxes, interquartile range; whiskers, 10th–90th percentiles. Inset, variance in disorder explained by family, by class and by shuffled families. **b**, Each row splits all genes by one label from UniProt or another database (for example, the UniProt keyword DNA-binding): genes that carry the label (filled) and all other genes (open). Dots, median percentage of residues in IDRs in each gene’s canonical isoform; orange, labelled genes more disordered than other genes; blue, less disordered; number of labelled genes in parentheses. Labels overlap, so a gene can count in many rows.

Single annotations gave a consistent picture when we compared the canonical disorder of genes carrying each UniProt label with that of all other genes (Fig. 3b). Genes annotated with DNA binding, transcription regulation, chromatin regulation, mRNA processing or developmental function were about twice as disordered as genes without them, as were genes with at least five phosphosites; genes with many interactors and genes in DisProt were about a quarter more disordered, and genes in condensates barely differed from other genes (median 24.7% versus 23.1%); transmembrane, secreted, mitochondrial and enzyme annotations marked the least disordered genes. A model with all annotations and protein length explained 31% of the variance in disorder (9,349 genes), against 20% for class and length. The class ordering did not depend on the predictor: MobiDB-lite consensus disorder from UniProt correlated with MetaPredict disorder per protein (*ρ* = 0.77) and ranked the classes similarly, although it called less disorder overall (Extended Data Fig. 2e).

Mode of disease inheritance also separated genes: genes with autosomal dominant disease were twice as disordered as genes with autosomal recessive disease (median 32% versus 16%), and their alternative exons and isoform-specific positions were more disordered relative to the rest of the gene (Extended Data Fig. 3).

### 2.3 Alternative exons encode more disorder than constitutive exons in every class

Across all genes, unique alternative exons encoded more disorder than constitutive exons (mean IDR fraction 0.380 versus 0.237). With genes as the unit of inference the difference was 0.143 (95% interval from resampling genes 0.134–0.152); it was 0.085 (0.075–0.095) after adjustment for exon length, exon position and number of transcripts, 0.174 within genes, 0.086 among internal exons only, and 0.108 and 0.073 when exon usage was defined on coding coordinates or across all GENCODE transcripts (Extended Data Fig. 2f). The global codon-level comparison gave the same answer at the level of residues: isoform-specific positions were more disordered than shared positions (41.2% versus 31.9%; difference 0.091, 0.073–0.108), and within genes the isoform-specific positions were the more disordered in 50% of genes and the less disordered in 35% (8,045 genes; Wilcoxon *P* = 1.2*×*10^*−*129^). The difference did not depend on how disorder was called: it was 0.083–0.159 with MetaPredict thresholds of 0.4–0.6, minimum IDR lengths of 10–50 residues or unsmoothed scores, 0.114 with MetaPredict V1 and 0.036 with MobiDB-lite on canonical isoforms, which calls far less disorder overall (6.7% of residues in the median protein, against 23.4%); its interval excluded zero in every case (Extended Data Fig. 2b).

The enrichment held in every protein class (Fig. 4a–c). The class-level difference ranged from 0.086 in secreted proteins and 0.089 in cytoskeletal proteins to 0.187 in other membrane proteins and 0.228 in receptors and channels, and its 95% interval excluded zero in all eleven classes. In metabolic enzymes, 23% of alternative exons encoded mostly disordered sequence against 9% of constitutive exons (odds ratio 2.95), so splicing introduces disorder even into the least disordered class. Annotations that marked membrane proteins widened the gap between alternative and constitutive exons: transmembrane (0.188 versus 0.129 in genes without the annotation), signal peptide, glycoprotein, cell membrane and receptor (Benjamini–Hochberg *q ≤* 0.002 for the interaction with exon usage); coiled coils weakened it (0.080 versus 0.156; *q <* 0.001). Classes also differed in how much of their exon content was alternative (Kruskal–Wallis *P* = 7.8 *×* 10^*−*43^; Fig. 4d), from a median of 39% of unique exons in kinases and phosphatases to 67% in DNA-binding transcription factors. Across the 50 families, the alternative–constitutive difference increased only weakly with family disorder (*ρ* = 0.29, *P* = 0.04), and some highly disordered families showed almost none, such as the SR splicing factors (0.03) and the C2H2 zinc-finger family (*−*0.01), against 0.29 in the paired homeobox and 0.20 in the ETS families.

**Fig. 4.**
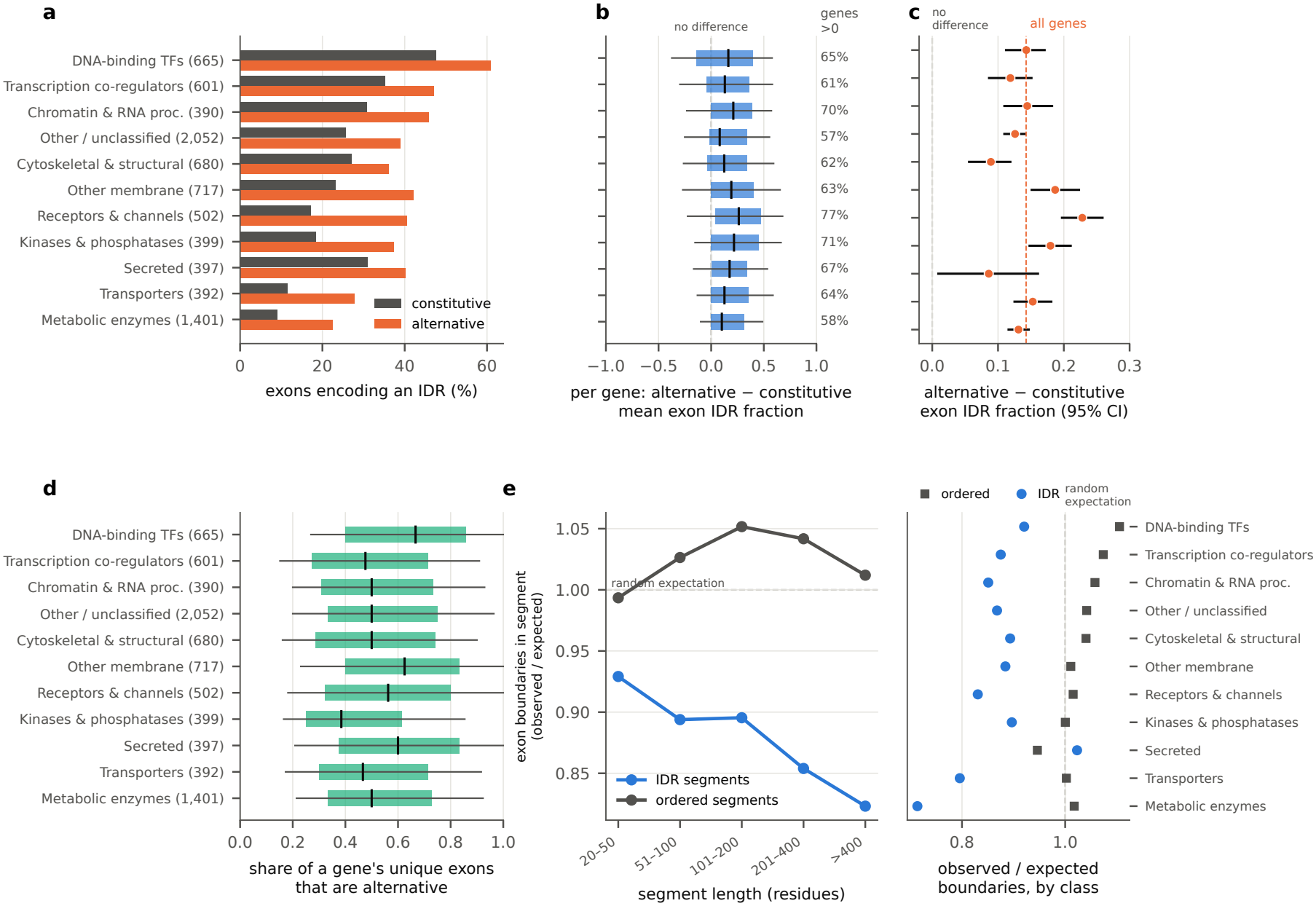
Alternative exons encode more disorder than constitutive exons in every protein class. **a**, Percentage of unique exons that are mostly disordered (IDR fraction *≥* 0.5), constitutive and alternative, per class; number of genes in parentheses. **b**, For each gene with both exon types, the alternative-minus-constitutive difference in mean exon IDR fraction; boxes, interquartile range; whiskers, 10th–90th percentiles; right, percentage of genes with a positive difference. **c**, Class-level difference with 95% intervals from 1,000 bootstrap resamples of genes; dashed line, all genes. **d**, Share of each gene’s unique exons that are alternative. **e**, Exon boundaries within IDR and ordered segments divided by the number expected for a segment of the same length placed uniformly at random in the same protein, by segment length (left) and by protein class (right).

IDRs contained fewer exon boundaries than segments of the same length placed at random in the same protein, by 7% at 20–50 residues and by 18% above 400 residues, whereas ordered segments contained as many as expected or slightly more (observed/expected 0.99–1.05; Fig. 4e). The depletion was present in ten of eleven classes, strongest in metabolic enzymes (0.71) and transporters (0.80), and absent in secreted proteins (1.02). IDRs are therefore modestly more exon-contained than their length predicts, not discrete exon modules.

### 2.4 Splicing changes IDRs mostly at their ends

Disorder increased with how rarely a position was used (Fig. 5a). Positions in every isoform were 32% disordered, positions in a majority or half of the isoforms 38–39% and positions in a minority of isoforms 51%. In 2,647 genes with at least three isoforms, minority positions were more disordered than majority positions within genes (0.51 versus 0.40; Wilcoxon *P* = 8.1 *×* 10^*−*39^).

**Fig. 5.**
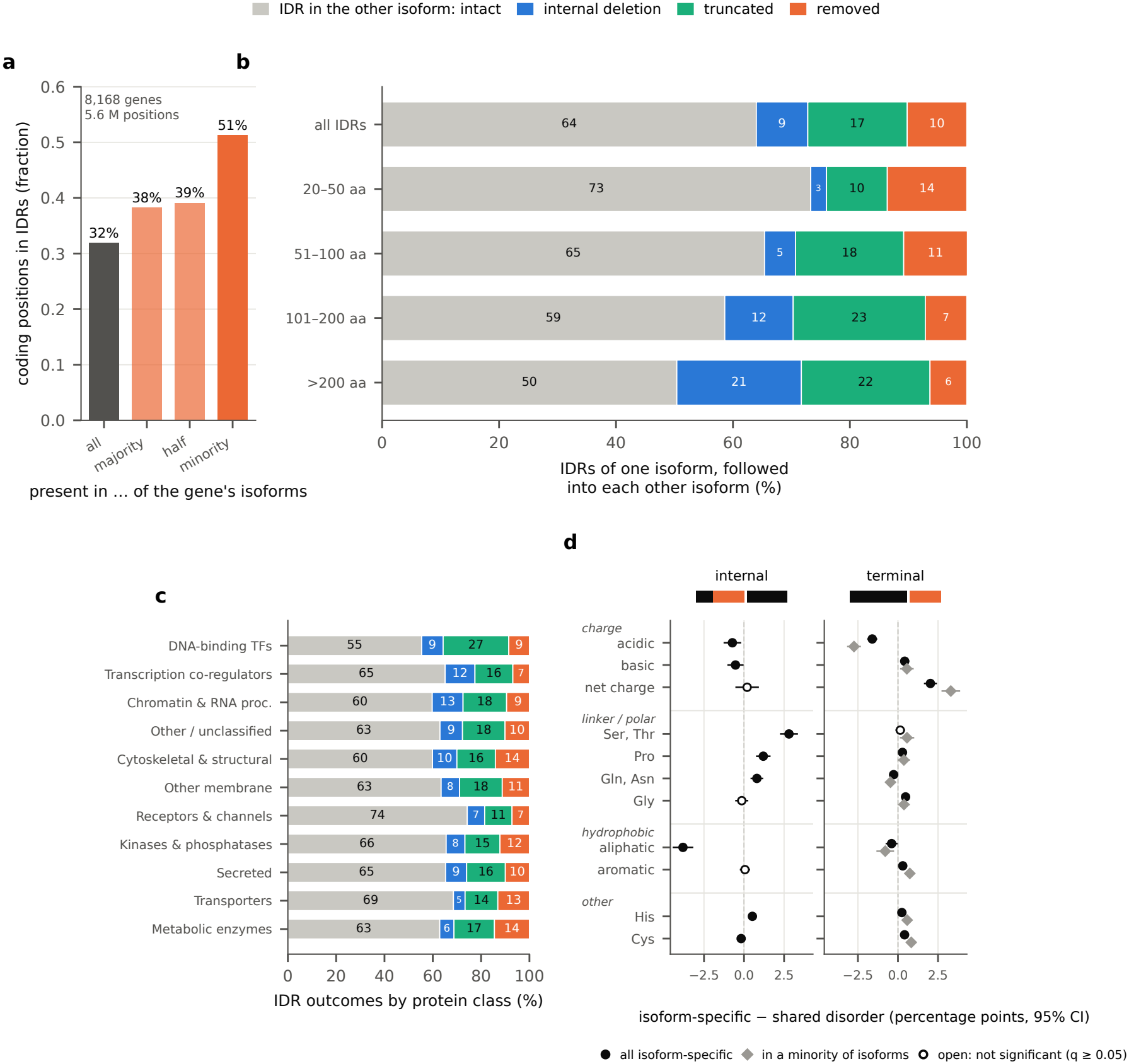
How splicing changes IDRs between isoforms, with residues shared only when the same genomic codon encodes them. **a**, Fraction of coding positions in IDRs by inclusion: present in all, a majority, half or a minority of a gene’s isoforms. **b**, Outcome of each IDR of one isoform in each other isoform of the gene (all ordered pairs), overall and by IDR length. **c**, IDR outcomes by protein class; the same IDRs classified by protein alignment are compared in Extended Data Fig. 4a. **d**, Composition of isoform-specific disorder minus the shared disorder of the same genes (gene-paired linear models; 95% intervals), for internal isoform-specific disorder (left; flanked by shared sequence, as in the icon) and terminal isoform-specific disorder (right; beyond the first or last shared position), all and in a minority of isoforms (grey diamonds). Rows are grouped into charge, linker/polar, hydrophobic and other residues. Filled, Benjamini–Hochberg *q <* 0.05; open, not significant. Net charge per residue is multiplied by 100. The full grid by inclusion is in Extended Data Fig. 6.

We followed each IDR of each isoform into every other isoform of its gene (98,994 IDR comparisons from 57,228 ordered isoform pairs in 7,195 genes; Fig. 5b). An IDR was intact in 64% of comparisons, truncated at one or both ends in 17%, removed entirely in 10% and internally deleted in 9%. Because the number of ordered pairs grows with the square of a gene’s isoform count, 5% of genes supplied 48% of these comparisons; with every gene weighted equally the outcomes were 60% intact (95% interval from resampling genes 59–61), 20% truncated, 14% removed and 6% internally deleted (Extended Data Fig. 4b). Internal deletion rose with isoform number, from 5% in genes with two isoforms to 12% in genes with six or more, and canonical IDRs followed into alternative isoforms were removed far more often (16%) than alternative IDRs followed into the canonical (3%), since canonical isoforms are mostly the longer (Extended Data Fig. 4c). Keeping an IDR’s codons kept its predicted disorder: for intact IDRs, 94% had at least 90% of their codons disordered in the other isoform’s own prediction (Extended Data Fig. 4d). The outcome depended on IDR length: IDRs of 20–50 residues were removed in 14% and internally deleted in 3%, whereas IDRs longer than 200 residues were internally deleted in 21% and removed in 6%. Classes differed in the same direction (Fig. 5c): DNA-binding transcription factors, with the longest IDRs, had the fewest intact IDRs (55%) and the most truncation (27%); metabolic enzymes and transporters had the most removal (14–15% and 13%); and receptors and channels had the most intact IDRs (74%).

These fractions differ from those obtained by aligning protein sequences in the usual way. For the same 23,849 canonical IDRs in alternative isoforms, both a longest-common-block matcher and a global Needleman–Wunsch alignment (BLOSUM62) called 22% internal deletions, against 10% at the codon level (Extended Data Fig. 4a). The excess came from isolated matches of one to four identical residues inside sequence that splicing had removed: discarding matched runs shorter than five residues gave 9.8% internal deletions with either method, and the alignment and codon classifications then agreed for 98% of IDRs (Extended Data Fig. 4a). Keying residues by codon reaches the same answer without a length threshold and while comparing every isoform with every other. Splicing therefore changes IDRs mainly at their ends, and internal deletion is common only in long IDRs. Two genes illustrate the modes (Extended Data Fig. 5): in PDLIM3, two disordered central exons of one isoform are replaced by a single different disordered exon in the other, and in MDM4 one isoform skips a fully disordered internal exon while another replaces all downstream exons with an alternative terminal exon.

The composition of isoform-specific disorder depended on where it lay (Fig. 5d; Extended Data Fig. 6). Compared with the shared disorder of the same genes, internal isoform-specific disorder (flanked by shared sequence; 1,454 segments) was depleted of aliphatic residues (*−*3.8 percentage points) and enriched in serine and threonine (+2.8), proline (+1.2), glutamine and asparagine (+0.8) and histidine (+0.5), with slightly fewer charged residues (*−*1.3) and no shift in net charge. Terminal isoform-specific disorder (4,214 segments) was instead depleted of acidic residues (*−*1.6) and more positively charged (net charge per residue +2.0), and modestly enriched in histidine, cysteine and aromatic residues. The terminal signature grew as inclusion fell: disorder at the ends of a minority of isoforms was depleted of acidic residues by 2.7 percentage points and shifted in net charge by +3.3, whereas disorder at the ends of most isoforms barely differed from shared disorder.

### 2.5 Disorder and splicing through gene age

Across the 10,450 genes with an age estimate, older genes encoded less disorder in all of their isoforms, canonical and alternative alike (32,349 isoforms; Spearman *ρ* = *−*0.19 between gene age and the gene’s mean disorder over its isoforms; median 28% of residues in isoforms of genes younger than 100 million years (Myr) and 15% in genes older than 1,000 Myr), as reported previously for canonical proteins[9]. The trend was negative in ten of eleven classes and significant in eight (Fig. 6a), strongest in transcription co-regulators (*ρ* = *−*0.28), cytoskeletal proteins and unclassified proteins (*−*0.27 each), and absent in kinases and phosphatases (*−*0.00, *P* = 0.94) and transporters (*−*0.07, *P* = 0.11). DNA-binding transcription factors were the one class in which disorder rose with gene age (*ρ* = +0.29, *P* = 2.7 *×* 10^*−*18^; median 29% in genes younger than 100 Myr and 66% in genes older than 1,000 Myr). The longest IDR of each isoform followed the same pattern (Fig. 6b): it was shorter in older genes overall (*ρ* = *−*0.12; median 85 residues below 100 Myr and 52 above 1,000 Myr), but in DNA-binding transcription factors it lengthened from 131 to 230 residues (*ρ* = +0.18).

**Fig. 6.**
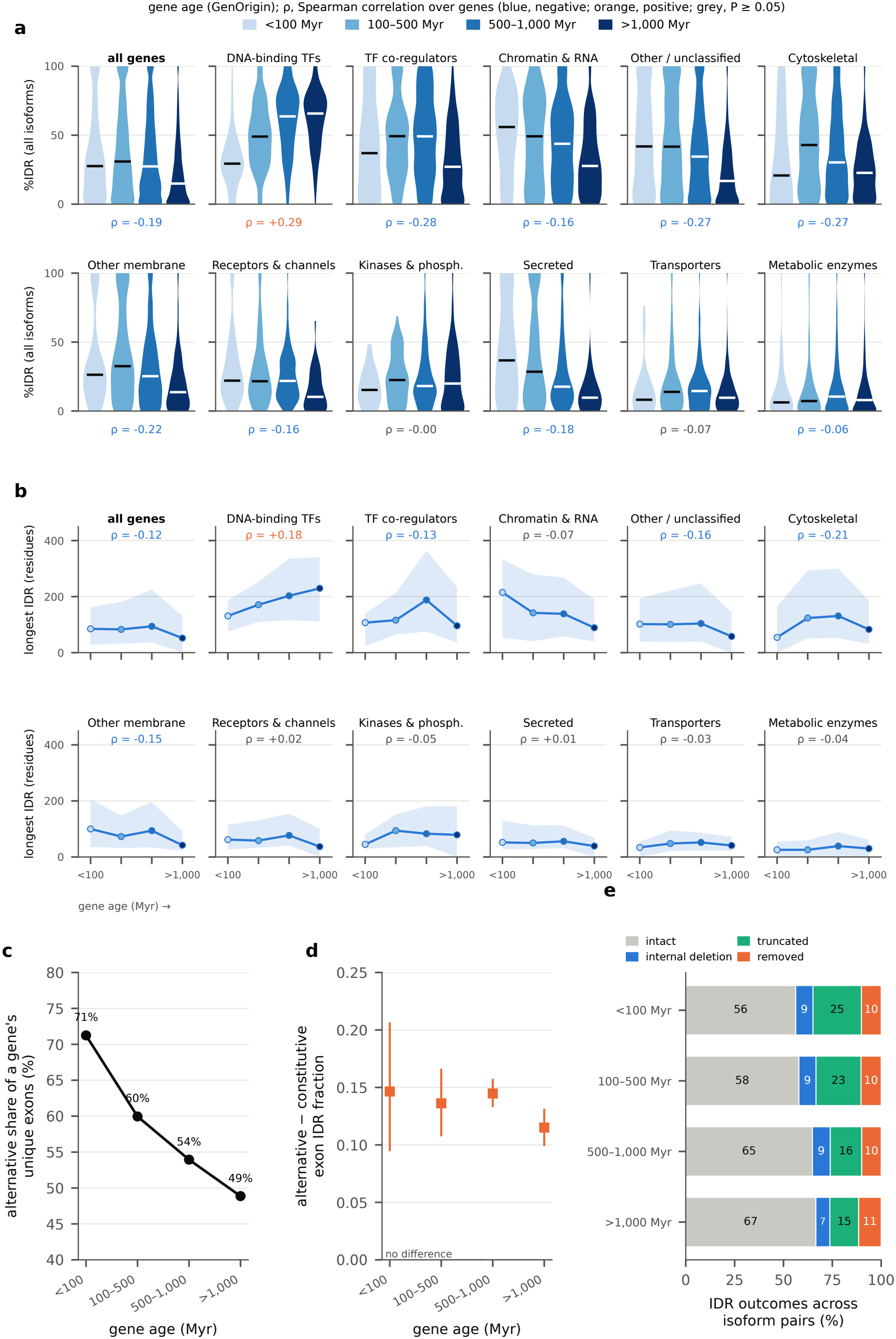
Disorder and splicing through gene age, by protein class. **a**, Percentage of residues in IDRs for every isoform, canonical and alternative, by gene age (GenOrigin), for all genes and each protein class; violins over isoforms, line at the median; *ρ*, Spearman correlation of gene age with each gene’s mean over its isoforms (blue, negative; orange, positive; grey, *P ≥* 0.05). **b**, Length of the longest IDR of every isoform by gene age (median, line; interquartile range, band; isoforms without an IDR count as zero). **c**, Share of each gene’s unique exons that are alternative, by gene age. **d**, Alternative-minus-constitutive difference in exon IDR fraction by gene age (95% intervals from resampling genes). **e**, IDR outcomes across all ordered isoform pairs, by gene 1a2ge.

Because families differ both in disorder and in age, we asked whether the age trend holds within families. It did not. Among 2,048 genes in 49 families with at least 15 dated genes, disorder fell by 0.23 percentage points per 100 Myr of gene age (95% interval 0.14–0.32), but with family fixed effects the slope was +0.04 (*−*0.05 to 0.13; *P* = 0.35). Across families, older families tended to be less disordered (*ρ* = *−*0.27 between family median age and median disorder, *P* = 0.06). Within individual families (Extended Data Fig. 7), the median correlation of canonical disorder with age was *−*0.07; no family had a significant negative correlation, and three had significant positive ones: the C2H2 zinc-finger (266 genes, *ρ* = 0.51), TRIM (43 genes, *ρ* = 0.50) and protein kinase families (344 genes, *ρ* = 0.17). Following every isoform through gene age within the 15 families that span at least three age bins gave the same picture (Extended Data Fig. 8): disorder and the longest IDR were flat in most families and rose with age in the C2H2 zinc-finger, TRIM, protein kinase and immunoglobulin families. The decline of disorder with gene age across the proteome therefore reflects which families are old rather than loss of disorder within families, and the positive trend in DNA-binding transcription factors coincides with the C2H2 zinc-finger family, whose young members are less disordered and make up most young transcription factors (92 of 111 younger than 100 Myr). Transcription factors outside that family showed no trend with age (*ρ* = *−*0.08, *P* = 0.05). No disorder definition gave a decline within families (slopes *−*0.04 to +0.09 per 100 Myr; Extended Data Fig. 2d), whereas the proteome-wide decline itself was weaker with MetaPredict V1 (*ρ* = *−*0.10) and MobiDB-lite (*−*0.06) than with MetaPredict v3 (*−*0.19).

Splicing also changed with gene age. Younger genes had a larger share of alternative exons (71% of unique exons in genes younger than 100 Myr, 60% at 100–500 Myr, 54% at 500–1,000 Myr and 49% in genes older than 1,000 Myr; Fig. 6c), although they did not have more isoforms (mean 2.7 versus 3.0). The alternative-minus-constitutive difference in exon disorder was positive at every age and somewhat smaller in the oldest genes (0.15 below 100 Myr and 0.12 above 1,000 Myr; Fig. 6d). IDR outcomes shifted most: in genes younger than 500 Myr, 23–25% of IDR comparisons were truncations and 56–58% intact, whereas in older genes 15–16% were truncations and 65–67% intact (Fig. 6e).

### 2.6 Exons with poorly conserved splice sites encode more, and different, disorder

Gene age compares different genes. To compare sequence of different ages within the same gene, we dated exons by the conservation of their splice sites: the number of 405 aligned mammalian genomes in which the GT donor or AG acceptor dinucleotide is conserved[21]. An internal exon is no older than its less conserved boundary, so we used the lower of its two counts; the count is a conservation score rather than a date, because it depends on how densely each clade is sampled. Of 90,817 internal coding exons with both splice sites matched, 1,842 (2.0%) had a splice site conserved in fewer than 100 genomes (poorly conserved), 6,705 (7.4%) in 100–349 and 82,270 (90.6%) in at least 350.

Exons with poorly conserved splice sites were more disordered than deeply conserved exons (mean IDR fraction 0.43 versus 0.25), more often alternative (72% versus 36%) and included in fewer of their gene’s transcripts (mean inclusion 0.43 versus 0.71; Fig. 7a). Within genes, adjusting for exon length, usage and transcript number, an exon with a poorly conserved splice site encoded 0.107 more IDR fraction than the gene’s other exons (0.087–0.127; *P* = 9.3*×*10^*−*26^; 7,653 genes; Extended Data Fig. 9b). The difference held among constitutive exons alone (0.094) and alternative exons alone (0.085), in nine of eleven classes, and in genes older than 1,000 Myr (0.093, 0.053–0.134), so old genes also carry young, disordered exons. Disordered sequence evolves quickly, and an exon that aligns poorly across species would also appear to lose its splice sites. Conservation of the exon’s own flanking bases is a poor control for this, because it falls with disorder even among exons whose splice sites are kept in at least 396 genomes (within genes, *−*5.4 genomes per unit IDR fraction, *P* = 4 *×* 10^*−*145^; Extended Data Fig. 2a); adjusting for it reverses the sign of the effect, and restricting to exons whose flanks are conserved in at least 350 genomes shrinks it to 0.038 (0.010–0.066; Extended Data Fig. 9). The 30 intronic bases next to each splice site measure whether the locus aligns without depending on the protein, and their conservation did not fall with disorder (+10.3 genomes). Adjusting for intronic conservation left the effect at 0.115 (0.095–0.134); it was 0.110 among exons whose intronic flanks align in at least 200 genomes, between 0.095 and 0.107 for thresholds of 50–200 genomes or for either splice site alone, and 0.090 (0.064–0.117) in length- and usage-matched pairs of exons within 1,021 genes (Extended Data Fig. 9b). With MetaPredict V1 the effect was 0.078 (0.059–0.097) and with MobiDB-lite on canonical exons 0.046 (0.026–0.066), each about the same fraction of the main estimate as for the alternative–constitutive difference (Extended Data Fig. 9b). Lining up exons by splice-site conservation within protein families gave the same picture (Extended Data Fig. 10). In 30 UniProt families with at least ten internal exons in each conservation bin, exons conserved in fewer than 350 genomes were more disordered than exons conserved in at least 380 in 25 families (median difference 0.11), with 95% intervals above zero in 14 and below zero in one (TRIM). The largest differences were in G-protein coupled receptors (0.45), cyclic nucleotide phosphodiesterases (0.36), ligand-gated ion channels (0.27) and protein kinases (0.25); the exceptions were mostly families whose exons are disordered at every conservation level, such as the SR splicing factors and the C2H2 zinc-finger family.

**Fig. 7.**
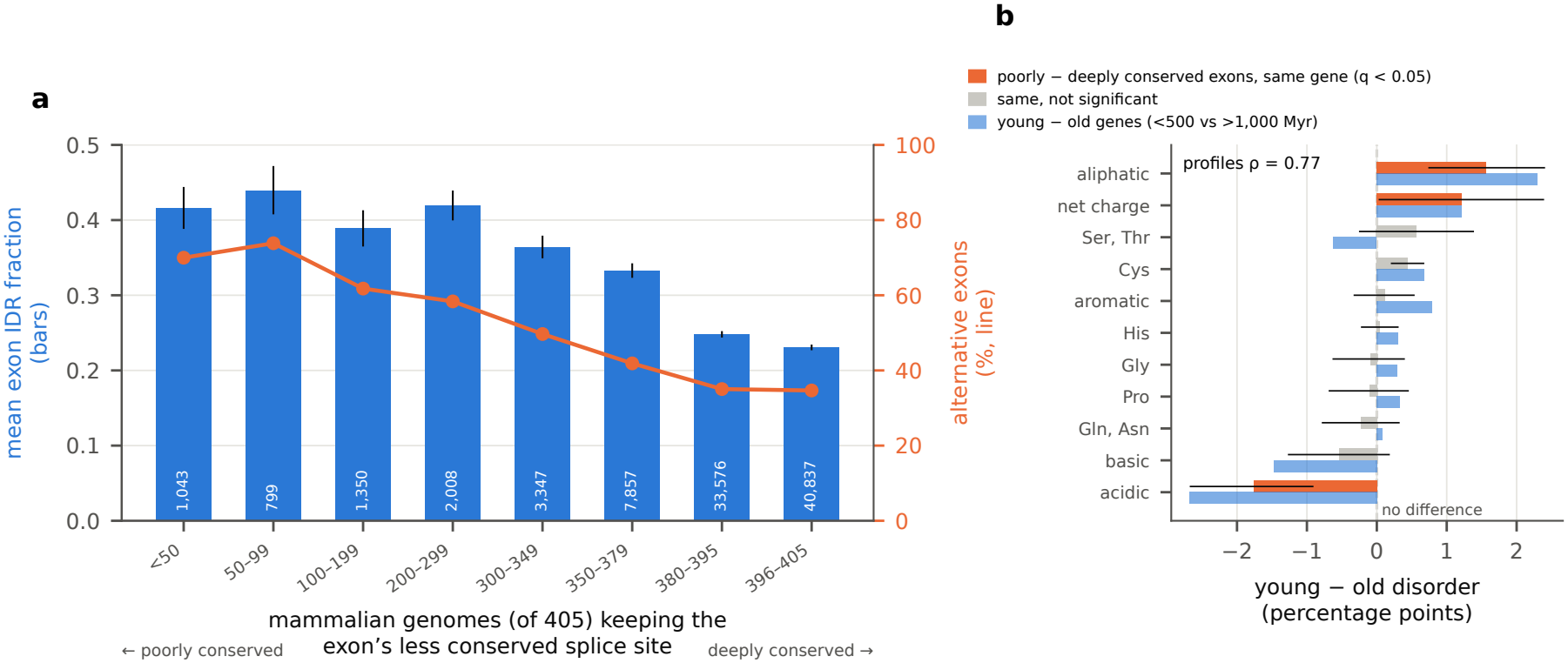
Exons with poorly conserved splice sites encode more, and different, disorder. **a**, Mean IDR fraction of internal coding exons (bars, 95% intervals; number of exons in white; left axis) and the share of them that are alternative (line; right axis), by the number of 405 mammalian genomes that keep the GT/AG of the exon’s less conserved splice site[21]. **b**, Composition of disordered residues from poorly conserved (*<*100) minus deeply conserved (*≥*350) exons of the same genes (lines, 95% intervals from resampling genes; orange, Benjamini–Hochberg *q <* 0.05; grey, not significant), beside the shared disorder of genes younger than 500 Myr minus genes older than 1,000 Myr (blue).

The composition of disorder from exons with poorly conserved splice sites differed from that of deeply conserved exons of the same genes (526 genes): it was depleted of acidic (*−*1.8 percentage points; *q* = 4 *×* 10^*−*7^) and charged residues (*−*2.3), more positively charged (net charge per residue +1.2) and enriched in aliphatic residues (+1.6), with no difference in aromatic content (Fig. 7b). The shared disorder of young genes (younger than 500 Myr) differed from that of old genes (older than 1,000 Myr) in the same direction (acidic *−*2.7, basic *−*1.5, aliphatic +2.3), and the two profiles were closely correlated across eleven features (*ρ* = 0.77, *P* = 0.005), but neither resembled internal isoform-specific disorder (*ρ* = 0.06). Young sequence is thus acid-poor whether it is a young gene or a young exon in an old gene.

## 3 Discussion

Alternative splicing is one of the main ways a single gene produces functionally distinct proteins, and most human multi-exon genes produce several isoforms[10, 11]. Because exons are discrete units of coding sequence, splicing can add, remove or replace whole segments of a protein, including the IDRs that carry short linear motifs, modification sites and condensate-forming sequence[4, 5]. Earlier proteome-wide studies established that alternatively spliced sequence is enriched for disorder[14, 15], but they left open whether this reflects a general property of proteins or the highly disordered regulatory proteins that dominate proteome-wide averages, what splicing does to an individual IDR, and how this relationship arose during evolution. Motivated by these questions, we mapped predicted disorder onto the coding exons and genomic codons of human protein isoforms and compared every isoform of a gene with every other, finding that splicing acts on disordered sequence in every kind of protein, that it remodels IDRs mostly at their ends, and that exons with poorly conserved splice sites contribute disordered sequence resembling that of young genes.

Notably, the enrichment of disorder in alternative exons is not confined to disordered regulators. It holds in each of the eleven protein classes, including metabolic enzymes and transporters, which are among the most ordered proteins, and it is strongest in receptors, channels and other membrane proteins (Fig. 4). One plausible explanation is that the folded transmembrane and extracellular domains of these proteins are encoded by constitutive exons, whereas splicing varies disordered cytoplasmic tails and loops that carry trafficking, signalling and scaffolding motifs. Consistent with this idea, isoforms of the same gene differ in disorder by a similar amount in every class (Fig. 2d), so splicing varies the disordered parts of mostly folded proteins as much as it varies proteins that are disordered throughout. Together, these observations suggest that splicing treats disorder as a modular layer that can be edited without disturbing the folded core of a protein.

How splicing changes an IDR depends on how isoforms are compared. When residues are matched by the genomic codon that encodes them, splicing most often changes an IDR at its ends, by truncating or removing it, and internal deletions become common only in IDRs longer than 200 residues (Fig. 5). This view fits the importance of alternative first and last exons as a source of protein diversity[22, 23], and the modest tendency of IDRs to lie within exons (Fig. 4e). Furthermore, the disordered sequence that splicing varies is not generic. Sequence inserted into or removed from the middle of a protein is enriched in serine, threonine and proline, residues characteristic of flexible, phosphorylatable linkers, whereas rarely used terminal sequence is depleted of acidic residues and more positively charged. One explanation is that the two kinds of splicing events act on different functional elements: internal events add or remove modification sites and linker length, whereas terminal events change the charge of a protein’s ends, which could alter its interactions, localization or partitioning into condensates.

Our results also refine the disorder-to-order model of protein evolution. Older genes encode less disorder in all of their isoforms, as reported for canonical proteins[9], but the relationship disap-pears when genes are compared within protein families (Fig. 6; Extended Data Figs 7 and 8). Several non-mutually exclusive explanations could contribute, including the origin of ancient families such as metabolic enzymes as ordered proteins, the origin of more recently arisen families as disordered ones, and differences among families in how long their members are retained. Although the available data cannot distinguish among these scenarios, they indicate that the proteome-wide trend largely reflects the composition of the gene repertoire rather than a gradual loss of disorder as individual proteins evolve. Consistent with the departure of transcription factors from the disorder-to-order expectation[9], DNA-binding transcription factors were the one class in which both disorder and the length of the longest IDR increased with gene age. This trend coincides with the recent expansion of the less disor-dered C2H2 zinc-finger family[24], and within the C2H2 zinc-finger, TRIM and protein kinase families disorder also rose with age, indicating that disorder can be gained or maintained over time in particular families.

Splice-site conservation connects these observations. Within genes, exons whose splice sites are conserved in few mammals encode more disorder than the gene’s other exons, are more often alternative and are included in fewer transcripts, and their disorder is depleted of acidic residues and more pos-itively charged, the same profile that separates the shared disorder of young and old genes (Fig. 7). One interpretation is that newly gained exons add disordered, acid-poor sequence to proteins of any age, consistent with the view that new coding sequence begins with a composition set more by its origin than by selection. However, a splice site conserved in few genomes may also have been lost in other species or misannotated[21], so the count measures how widely a splice signal is shared rather than when an exon arose. The association holds when the alignability of the surrounding introns is controlled for, in matched exon pairs and with other disorder predictors, although its size depends on how disorder is defined. If per-species splice-site presence confirms that these exons were recently gained, they would represent one route by which disordered sequence enters proteins.

Extending to human disease, genes associated with dominant disease were more disordered than genes associated with recessive disease, and their alternative exons were more disordered relative to their constitutive exons (Extended Data Fig. 3). This pattern is consistent with dosage sensitivity and with gain-of-function or dominant-negative mechanisms in disordered proteins, and suggests that vari-ants affecting which disordered exons are included could have dominant effects, although the association is correlational and confounded with protein function.

Our findings suggest that the role of splicing with respect to intrinsic disorder may not simply be to switch disordered regions on or off, but to edit them. Unlike folded domains, which are mostly encoded by constitutive exons and shared across a gene’s isoforms, IDRs appear to be where isoforms of a gene differ: at their ends, in their composition and in sequence that is evolutionarily young. This interpretation provides a unifying explanation for why alternative exons are disordered in every protein class, why splicing changes IDRs mostly at their ends, and why young exons resemble young genes. Rather than being incidental to splicing, IDRs emerge as the parts of proteins through which splicing most readily diversifies protein function.

There are several limitations of this work. For instance, although the predictor we used performs well in independent assessments[18, 25] and our main results hold with an earlier version of it and with MobiDB-lite, disorder is predicted, and future studies will ideally leverage experimentally vali-dated IDRs. MetaPredict v3 is trained partly on AlphaFold2 confidence, which is low for sequences with few homologues, so absolute levels of disorder differ between predictors. Furthermore, the binary classification of a segment as disordered or ordered does not capture folding upon binding or other context-dependent transitions. Exon usage reflects annotated transcripts rather than measured inclu-sion, so some rarely used exons may belong to minor or non-functional isoforms[22, 23]. Finally, gene ages date the origin of a gene rather than of its exons, the protein classes are rule-based and mutually exclusive, and all associations are correlational. Future work integrating isoform-resolved expression and splicing quantitative trait loci[26, 27], per-species splice-site presence and population-genetic con-straint on isoform-specific and shared disorder would test whether the disordered sequence that splicing adds is used and selected.

In conclusion, alternative splicing acts on intrinsic disorder in every class of human protein, editing IDRs mostly at their ends and adding compositionally distinct disordered sequence. Our findings demonstrate that the evolutionary and functional consequences of disorder are best understood at the level of isoforms and protein families rather than proteome-wide averages. More broadly, our findings raise the possibility that splicing has provided a recurrent route by which disordered sequence, and the regulatory interactions it encodes, enters and diversifies human proteins.

## 4 Methods

### 4.1 Protein isoform cohort

Reviewed human UniProtKB/Swiss-Prot entries including isoforms (query reviewed:true AND model organism:9606; 42,562 sequences, of which 20,431 canonical and 22,131 isoforms) from UniProt release 2026 01 (published 28 January 2026; downloaded 2 June 2026) were filtered to entries with a named gene, a length of at least 20 residues and protein-existence level below 5, and restricted to genes with at least two isoforms[28] (parameter TFIDR MIN ISOFORMS; the earlier cohort of genes with at least three isoforms is compared in Supplementary Table 1).

### 4.2 Disorder prediction

Disorder was predicted with MetaPredict (v3.1)[18], whose disordered-domain call smooths per-residue scores, uses a threshold of 0.5, closes short gaps and has a minimum IDR size of 12 residues; disordered domains of at least 20 residues were retained as IDRs, and %IDR is the percentage of residues in retained IDRs. Re-running MetaPredict 3.1.01 on 400 random isoforms reproduced the stored IDRs exactly. As an independent measure, we parsed MobiDB-lite consensus disordered regions from the UniProt ‘Region’ annotations of each canonical entry.

### 4.3 Transcript mapping and genomic projection

Isoform-tagged Ensembl cross-references from UniProt nominated candidate transcripts. Each candi-date’s GENCODE v50 (GRCh38) translation was compared with the UniProt sequence and only exact matches were used[29]. Isoforms with one coding structure (24,803) were projected to GRCh38 through a strand-aware array of coding positions, and every projected segment was checked by translation (all 76,781 passed). Exon IDR fraction is the fraction of an exon’s encoded residues in retained IDRs; an exon with IDR fraction *≥* 0.5 is mostly disordered.

### 4.4 Exon usage and statistics

For genes with at least two mapped transcripts, an exon (Ensembl identifier) was alternative if it occurs in some but not all of the gene’s mapped transcripts and constitutive otherwise. Unique exons are (gene, exon) pairs. Genes were the unit of inference: 95% intervals from resampling genes with replacement (1,000 resamples) of the alternative-minus-constitutive difference in mean exon IDR fraction; linear and logistic models with log_10_ exon length, exon position and log_2_ number of transcripts and gene-clustered errors; gene fixed effects; and paired Wilcoxon tests. Sensitivity analyses keyed exons by coding coordinates, used all GENCODE protein-coding transcripts, or used the inclusion fraction.

### 4.5 Global comparison of isoforms by genomic codon

Each residue of each mapped isoform was keyed by the genomic coordinate of the first base of its codon. For each gene, a position’s inclusion is the fraction of the gene’s mapped isoforms that encode it: shared (all), majority (*>*0.5), half (0.5) or minority (*<*0.5). A position was disordered if it lay in an IDR in at least half of the isoforms that encode it. Isoform-specific positions were internal if shared positions lay on both sides in the gene’s coordinate order and terminal otherwise. For IDR outcomes, each IDR of isoform *i* was followed into each other isoform *j* of the gene (all ordered pairs): intact if every codon of the IDR is in *j*, removed if none is, truncated if the missing codons form runs only at one or both ends of the IDR, and internally deleted if any missing run is internal. Outcomes were summarised pooled over comparisons and with each gene weighted equally (95% intervals from 300 resamples of genes), by the number of isoforms in the gene and by direction (canonical to alternative, alternative to canonical, alternative to alternative). A codon spanning an exon junction is keyed by its first base, so a codon whose second and third bases come from a different exon in the other isoform (a shifted splice site, or an insertion in the other isoform) counts as shared; keying codons by all three bases instead changed 4.1% of comparisons, almost all from intact to a one-residue internal deletion. To compare with protein alignment, canonical IDRs were also classified in each alternative isoform from difflib.SequenceMatcher matching blocks (with and without its junk heuristic, and keeping blocks of at least five or ten residues) and from global Needleman–Wunsch alignments (Biopython PairwiseAligner, BLOSUM62, gap open *−*10, extend *−*0.5; identical aligned residues counted as shared, with or without a minimum run of five), with the same definitions applied to shared residues; ten pairs longer than 6,000 residues were not aligned. For each IDR kept in another isoform, we recorded the fraction of its kept codons inside an IDR of that isoform’s own prediction.

### 4.6 Composition of isoform-specific disorder

Disordered positions were grouped into contiguous segments by inclusion class and position (shared; majority, half or minority; internal or terminal), keeping segments of at least ten residues. For each segment we computed the fractions of aromatic (F, W, Y), acidic (D, E), basic (K, R), histidine, cysteine, proline, serine and threonine, glutamine and asparagine, glycine and aliphatic (A, I, L, M, V) residues, the fraction of charged residues and the net charge per residue. Each isoform-specific group was compared with the shared disorder of the same genes with linear models adjusted for log_10_ segment size and gene-clustered errors, and P values were corrected across features within each group (Benjamini–Hochberg).

### 4.7 Length-controlled exon modularity

For each IDR and ordered segment of a mapped isoform, internal coding-exon boundaries within the segment were counted and compared with the exact expected count for a segment of the same length placed uniformly at random in the same protein.

### 4.8 Protein classes, families and annotations

Gene-level annotations were taken from UniProt (keywords, ‘Protein families’, EC numbers, subcellular location, transmembrane and signal-peptide features, modified residues, coiled coils, interactions, PDB cross-references and ‘Involvement in disease’), from DisProt and from CD-CODE condensate member-ship. Classes were assigned in the order listed in Results; a gene takes the first class whose rule it meets: DNA-binding transcription factor (HumanTFs[19]), transcription co-regulator (keyword Transcription regulation or DNA-binding), chromatin and RNA processing (keywords Chromatin regulator, mRNA processing or RNA-binding), kinase or phosphatase (keywords Kinase or Protein phosphatase), recep-tor or channel (keywords Receptor, G-protein coupled receptor or Ion channel), transporter (keyword Transport with a transmembrane segment), metabolic enzyme (EC number), cytoskeletal or structural (keyword Cytoskeleton or cytoskeletal location), secreted (signal peptide without a transmembrane segment), other membrane (transmembrane segment) and other. Family is the first UniProt family assignment; families with at least 15 genes were analysed, and variance explained (*η*^2^) was compared with 1,000 permutations of family labels. For each annotation, disorder of genes with and without it was compared with Mann–Whitney tests, and its effect on the alternative–constitutive difference was tested as the interaction of usage and annotation in the clustered exon model, with Benjamini–Hochberg correction.

### 4.9 Gene age and evolutionary analyses

Gene ages (million years) are from GenOrigin[20] and were binned as *<*100, 100–500, 500–1,000 and *>*1,000 Myr. For each class, we computed Spearman correlations of canonical disorder with age and regressed gene age on disorder (ordinary least squares, heteroscedasticity-robust errors), as in ref.[9]. To separate trends within and between families, canonical %IDR was regressed on gene age (per 100 Myr) among genes in UniProt families with at least 15 dated genes, without and with family fixed effects, and Spearman correlations of disorder with age were computed within each family with genes in at least two age bins (Benjamini–Hochberg correction across families). The same within-family and outside-family comparison was made for the C2H2 zinc-finger family among DNA-binding transcription factors. The alternative–constitutive difference, isoform-specific minus shared disorder and IDR outcomes were computed within each age bin as above. Composition of shared disorder was compared between genes younger than 500 Myr and older than 1,000 Myr with linear models adjusted for log_10_ segment size, and across all four bins with Kruskal–Wallis tests. Modes of inheritance were parsed from UniProt ‘Involvement in disease’ text (autosomal dominant, autosomal recessive or both); the dominant-recessive difference in disorder was adjusted for class and log_10_ length. For every isoform, canonical and alternative, we computed the percentage of residues in IDRs, the length of its longest IDR (zero without an IDR) and its mean IDR length; trends with gene age were tested with Spearman correlations on each gene’s mean over its isoforms, so that genes with many isoforms do not dominate, for all genes, each protein class and each UniProt family with at least 15 dated genes.

### 4.10 Exon conservation

Splice-site conservation was taken from ref.[21]: for each annotated splice site (GENCODE v45, RefSeq 110, CHESS 3 and MANE), the number of 405 genomes in the UCSC 470-mammal alignment in which the GT (donor) or AG (acceptor) dinucleotide is conserved, together with per-position conservation of the 30 bases on each side. Decoy sites were excluded and each site took its maximum count across annotations. Coding exons were matched by GRCh38 coordinate to their acceptor (exon start *−*2 on the plus strand, end +2 on the minus strand) and donor (end +1 on the plus strand, start *−*1 on the minus strand); 91% of acceptors and 89% of donors matched, the remainder being first or last exons or sites absent from GENCODE v45. Internal exons with both sites took the lower count. Exon IDR fraction was modelled on an indicator of an exon with a poorly conserved splice site (*<*100 genomes) with gene fixed effects, log_10_ exon length, usage and log_2_ number of transcripts, and gene-clustered errors. Alignability was measured from the per-position conservation of the 30 bases on each side of each splice site: the median over the 30 exonic bases (the lower of the exon’s two sites), and the median over the 30 intronic bases of the less conserved site. Because exonic conservation reflects the evolutionary rate of the encoded protein, which is faster in disordered sequence, we used intronic conservation as the alignability covariate and threshold. Exons with poorly conserved splice sites were also matched to an exon of the same gene and usage within a factor of 1.5 in length, and the difference in IDR fraction was averaged per gene and bootstrapped over genes (1,000 resamples). Within each UniProt family with at least 15 genes and at least ten internal exons in each of four conservation bins (*<*350, 350–379, 380–395 and *≥*396 genomes), mean exon IDR fraction was computed per bin, and the difference between exons conserved in fewer than 350 and at least 380 genomes was bootstrapped over genes (500 resamples). Composition of disordered residues was compared between poorly conserved and deeply conserved (*≥*350) exons within genes with at least ten disordered residues in each set (gene-paired Wilcoxon tests, 95% intervals from resampling genes, Benjamini–Hochberg correction).

### 4.11 Sensitivity to the disorder definition

The main results were recomputed with other disorder definitions: MetaPredict v3 disordered domains at thresholds 0.4 and 0.6 and with minimum IDR lengths of 10, 30 and 50 residues, computed from stored per-residue scores; runs of at least 20 residues with raw v3 scores *≥* 0.5 without smoothing; MetaPredict V1, the earlier version of the predictor (disordered domains at its threshold of 0.3, and raw runs); and MobiDB-lite consensus ‘Disordered’ regions from UniProt, available for canonical isoforms only. For each call we recomputed the class medians, the alternative-minus-constitutive difference in exon IDR fraction, the within-gene effect of an exon with a poorly conserved splice site and the change in disorder with gene age within families. Non-standard residues (U, B, Z, X) were replaced by C, N, Q and A for prediction.

## Acknowledgements

We thank members of the Akey laboratory for helpful feedback and comments related to this work. This work was supported by NIH grant R35GM161407 (JMA).

## Declarations

- **Funding:** NIH grant R35GM161407 (JMA).
- **Competing interests:** The authors declare no competing interests.
- **Ethics approval and consent to participate:** Not applicable.
- **Consent for publication:** Not applicable.
- **Data availability:** All input data are public: UniProtKB/Swiss-Prot, GENCODE v50, HumanTFs, GenOrigin, DisProt, CD-CODE and the splice-site conservation tables of Minkin and Salzberg. Derived tables are available from the project repository.
- **Code availability:** Analysis code is available at https://github.com/ph-an/as-idr-isoforms.

## Extended Data

**Extended Data Table 1:**
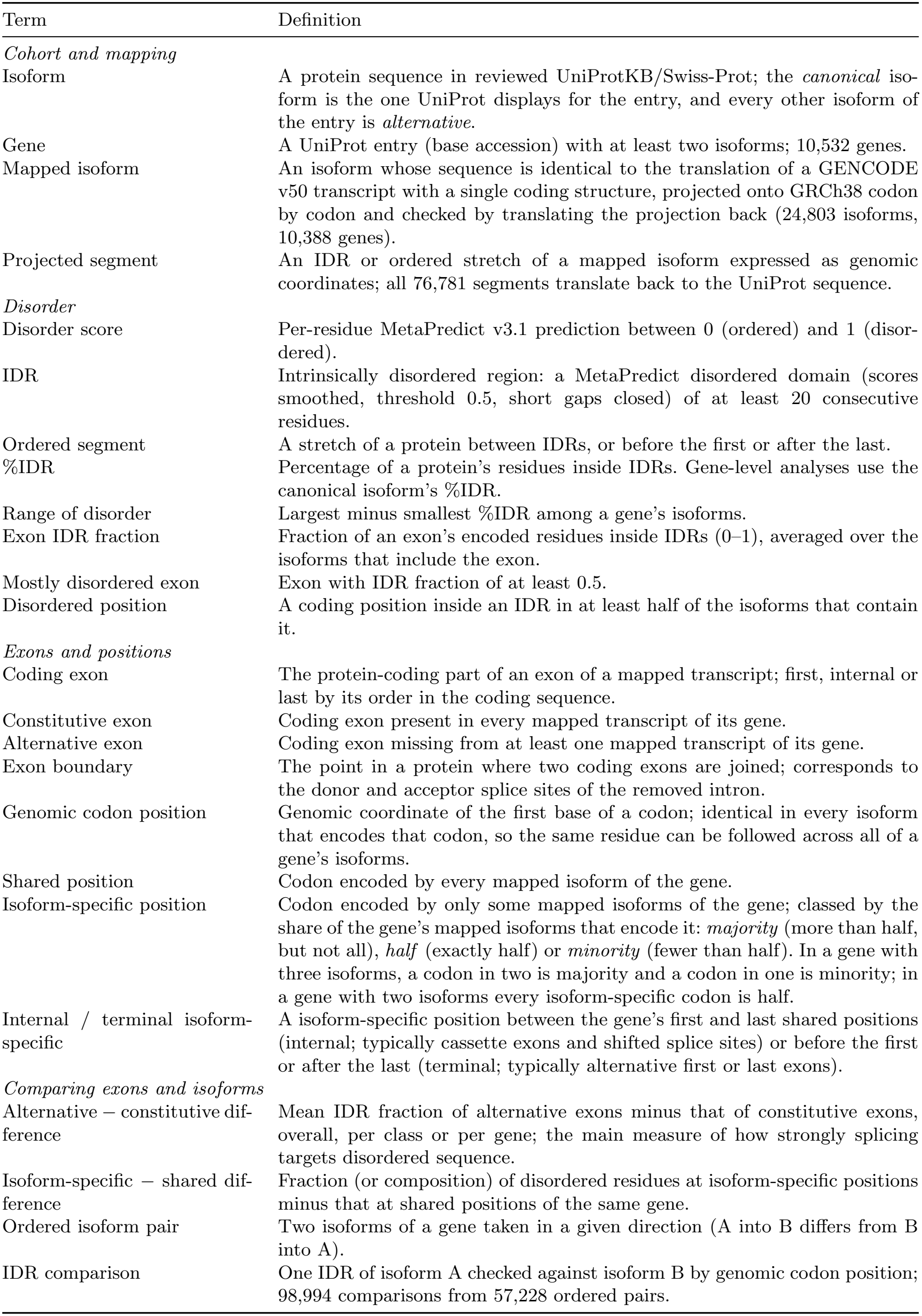
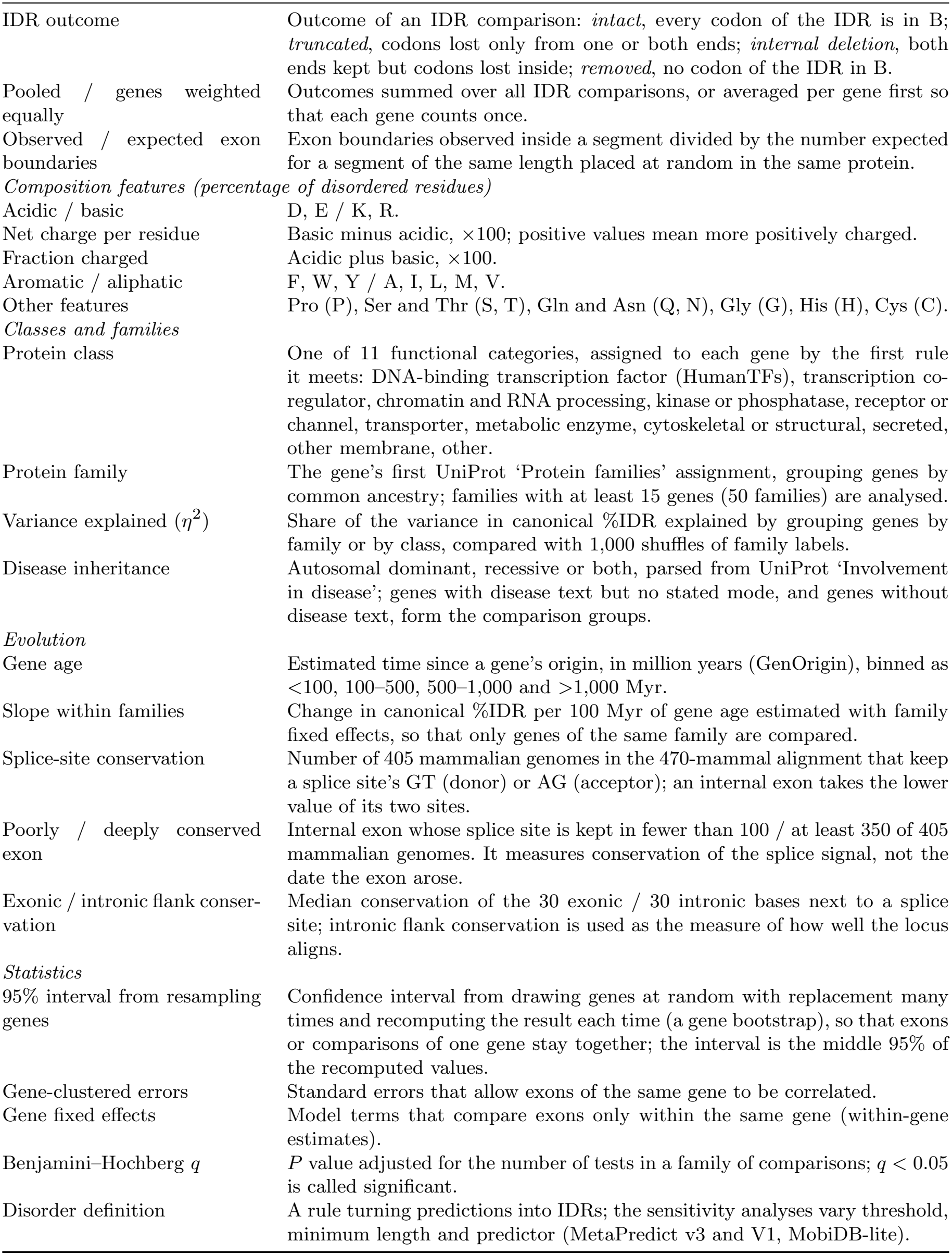
Glossary of terms and metrics used in the analyses.

**Extended Data Fig. 1.**
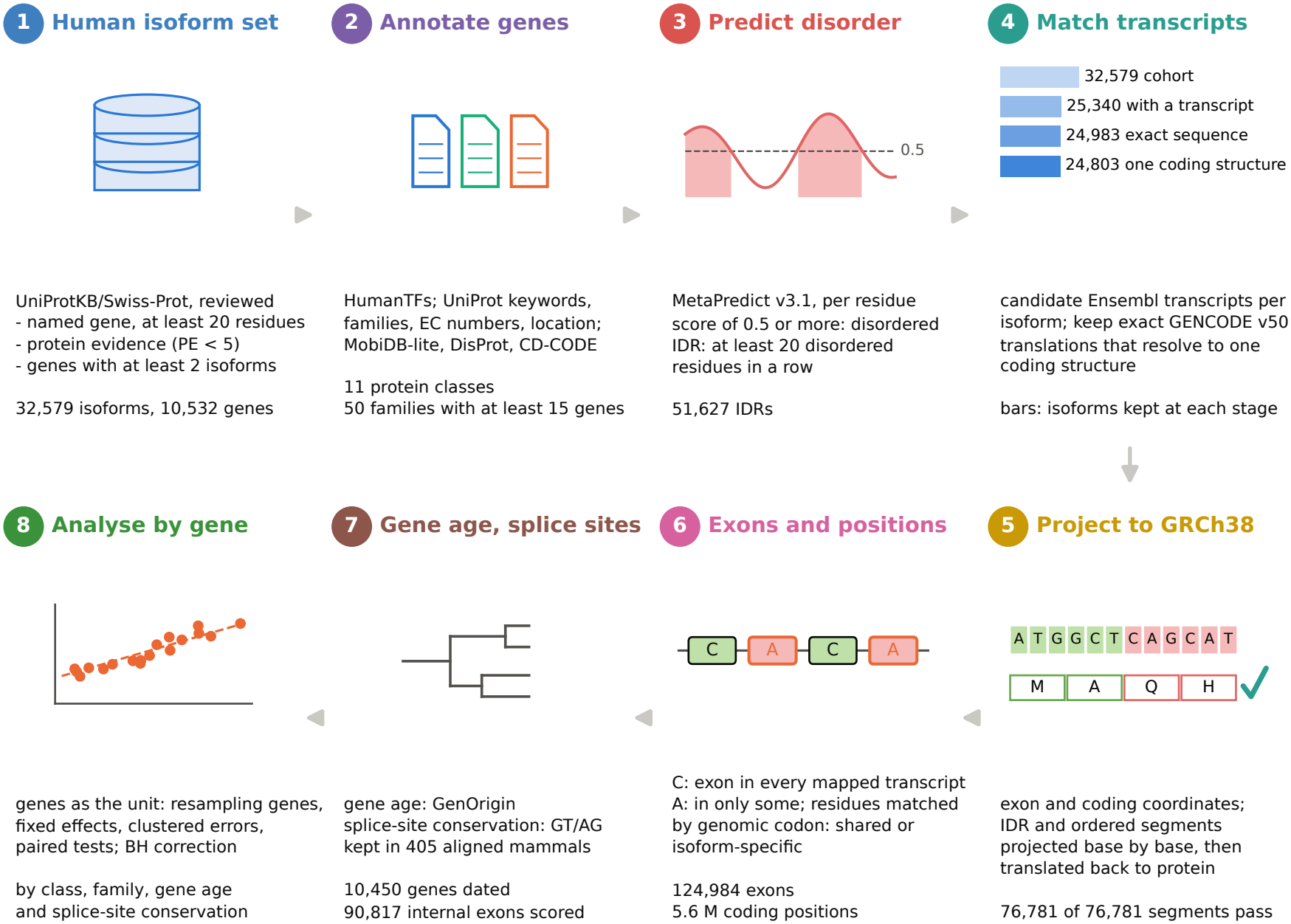
Methods workflow. Eight steps from reviewed UniProt isoforms to the analyses, with the number of isoforms, IDRs, genes, exons or positions at each step. In step 4, bars show the isoforms kept at each transcript-matching stage. Redrawn and updated from the 2026 QCB SURP poster.

**Extended Data Fig. 2.**
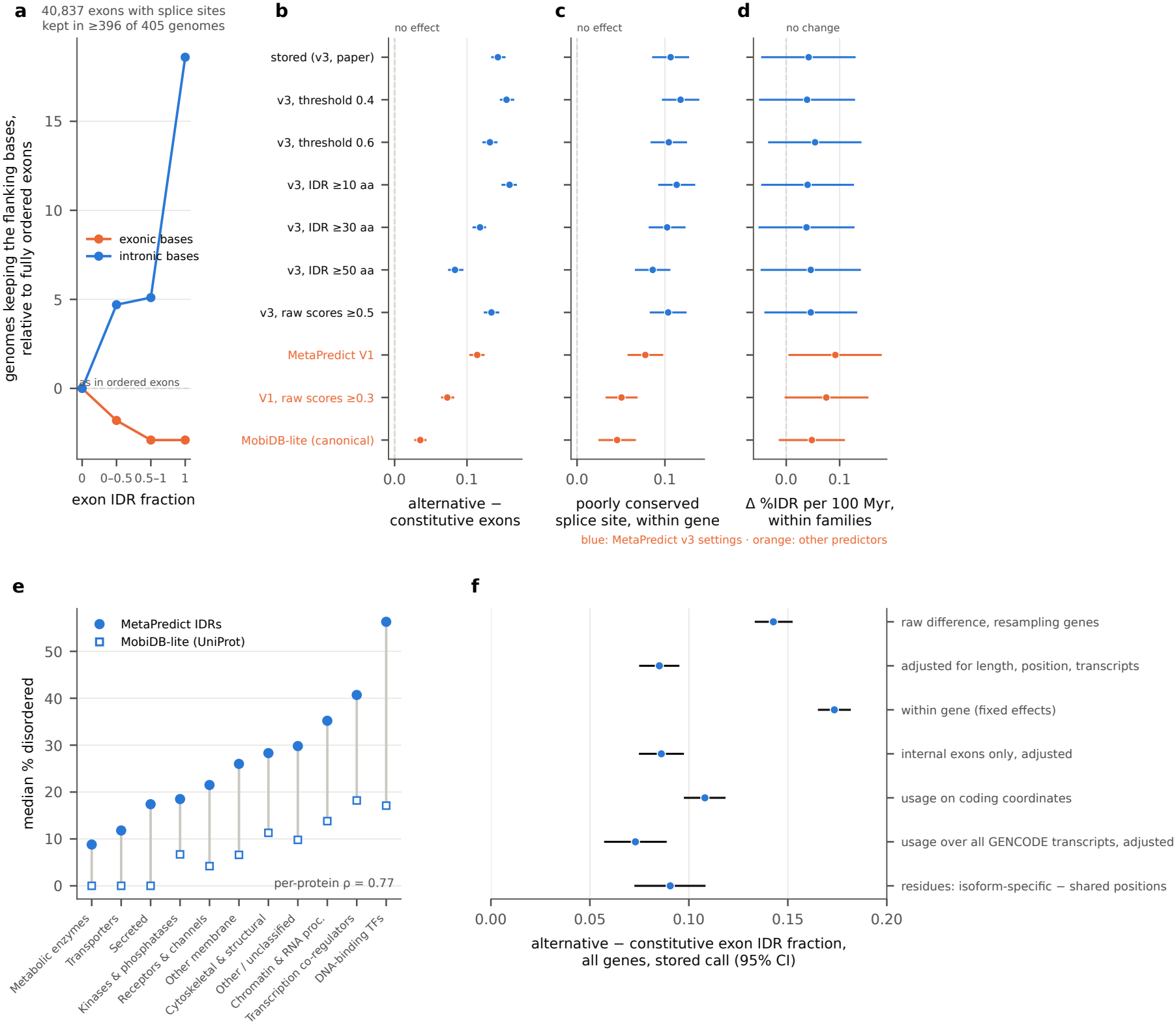
Robustness of the main results. **a**, Among internal exons whose splice sites are kept in at least 396 of 405 genomes, conservation of the 30 exonic and the 30 intronic bases next to the splice sites, by exon IDR fraction, relative to fully ordered exons. **b–d**, The alternative-minus-constitutive difference in exon IDR fraction (95% interval from resampling genes), the within-gene effect of an exon with a poorly conserved splice site (gene fixed effects) and the change in canonical %IDR per 100 Myr of gene age within families (family fixed effects) under each disorder definition (Methods); bars, 95% intervals; blue, MetaPredict v3 settings; orange, other predictors. **e**, Median disorder per class from MetaPredict IDRs and from MobiDB-lite consensus disorder in UniProt. **f**, The all-gene alternative-minus-constitutive difference in exon IDR fraction under different estimators and definitions of usage: raw difference with a 95% interval from resampling genes; linear model adjusted for log exon length, exon position and log number of transcripts with gene-clustered errors; within-gene estimate; adjusted model on internal exons; usage on coding coordinates; usage over all GENCODE protein-coding transcripts; and isoform-specific minus shared positions in the global codon-level comparison. Bars, 95% intervals.

**Extended Data Fig. 3.**
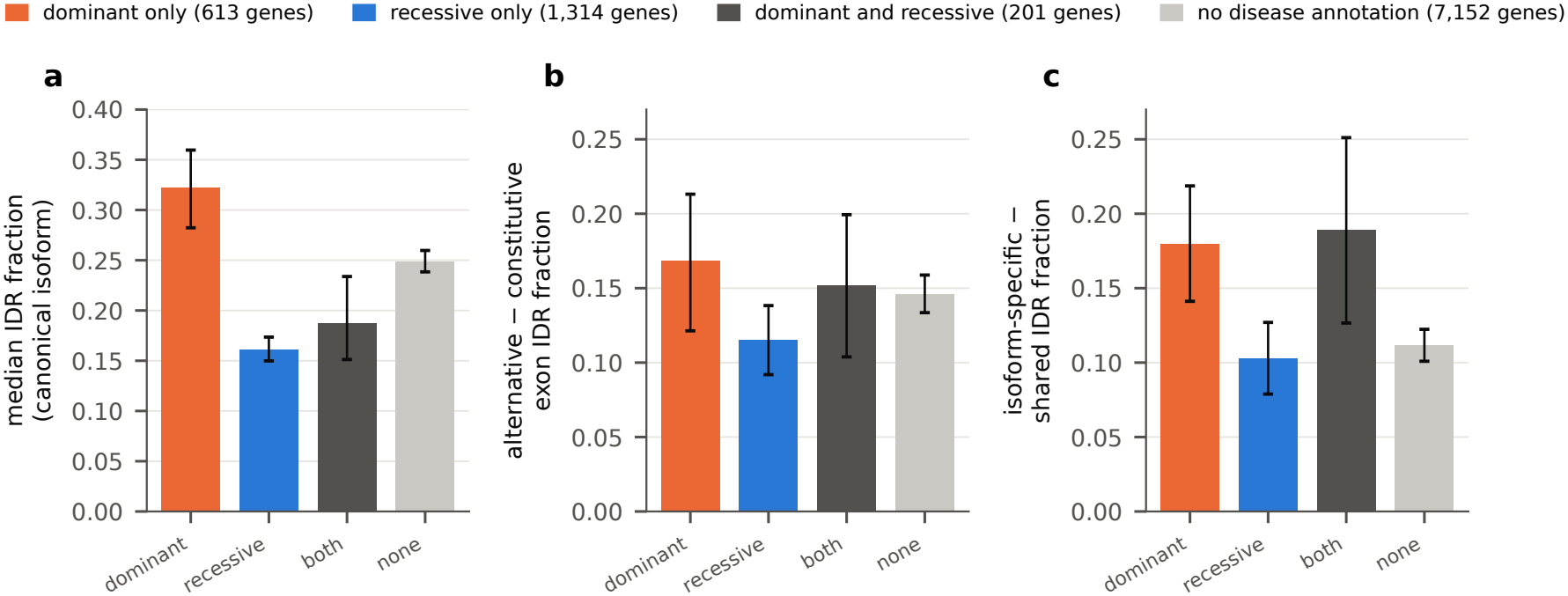
Disease inheritance and disorder. Genes with autosomal dominant, autosomal recessive or both modes of disease inheritance in UniProt (‘Involvement in disease’), and genes with no disease annotation (7,152); 1,252 genes with disease text that states neither mode are not shown. Bars, 95% intervals from resampling genes. Genes with dominant disease were more disordered than genes with recessive disease (Mann–Whitney *P* = 1.8 *×* 10^*−*29^; +6.0 percentage points, 3.8–8.3, after adjustment for class and length) and had more isoforms (mean 3.7 versus 3.1). **a**, Median IDR fraction of the canonical isoform. **b**, Alternative-minus-constitutive difference in exon IDR fraction. **c**, Isoform-specific minus shared difference in IDR fraction.

**Extended Data Fig. 4.**
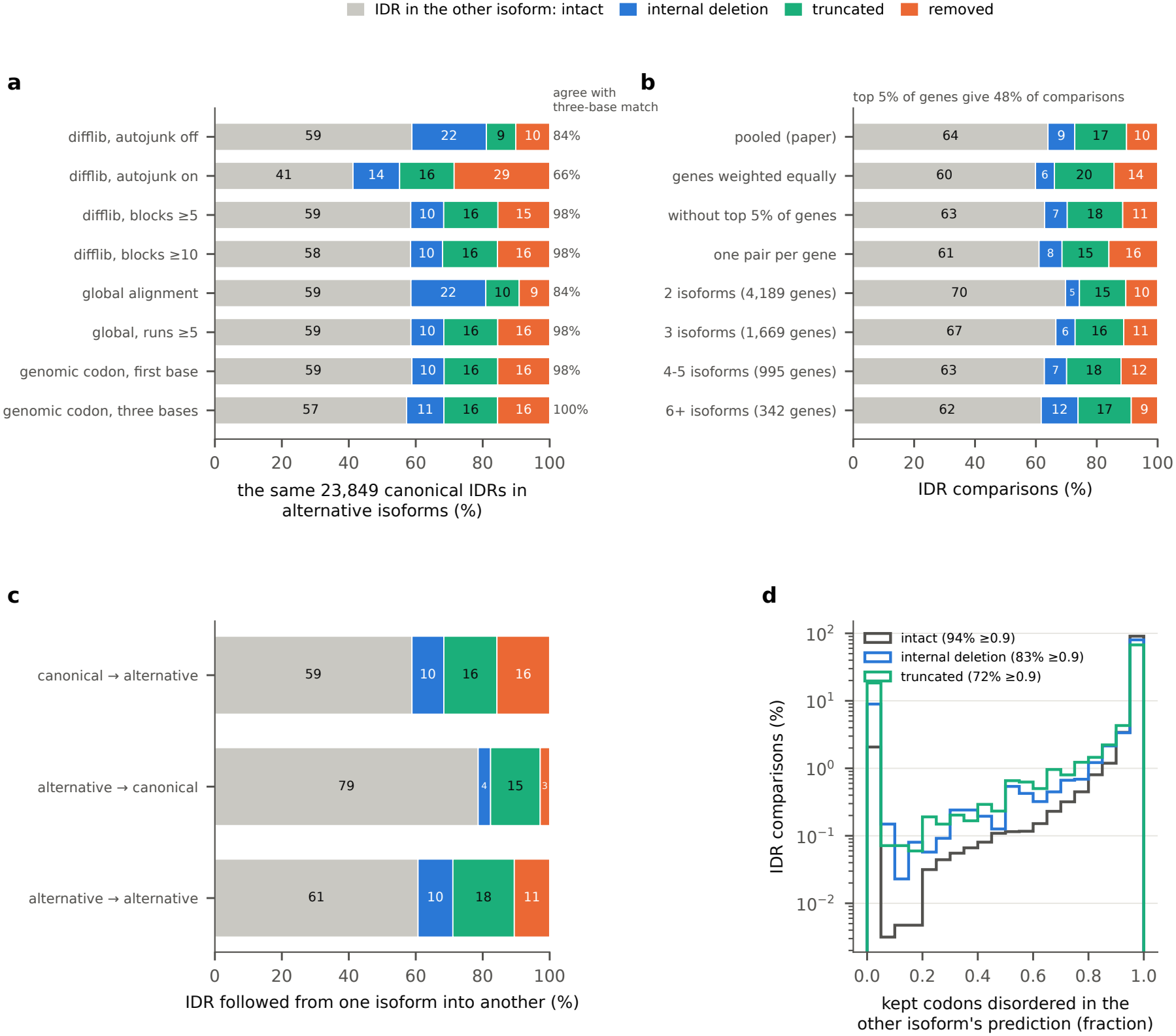
Validation of the codon-level IDR outcomes. **a**, The same canonical IDRs in alternative isoforms classified by difflib matching blocks (junk heuristic off or on; blocks of at least 5 or 10 residues), by global Needleman–Wunsch alignment (all identical aligned residues, or identical runs of at least five), and by genomic codon position (first base, as used throughout, or all three bases); right, agreement with the three-base genomic codon position. **b**, Outcomes over all ordered isoform pairs: pooled, with genes weighted equally, without the 5% of genes contributing most comparisons, with one random canonical–alternative pair per gene, and by the number of isoforms in the gene. **c**, Outcomes by direction. **d**, Fraction of an IDR’s kept codons that lie in an IDR of the other isoform’s own prediction, by outcome (log scale).

**Extended Data Fig. 5.**
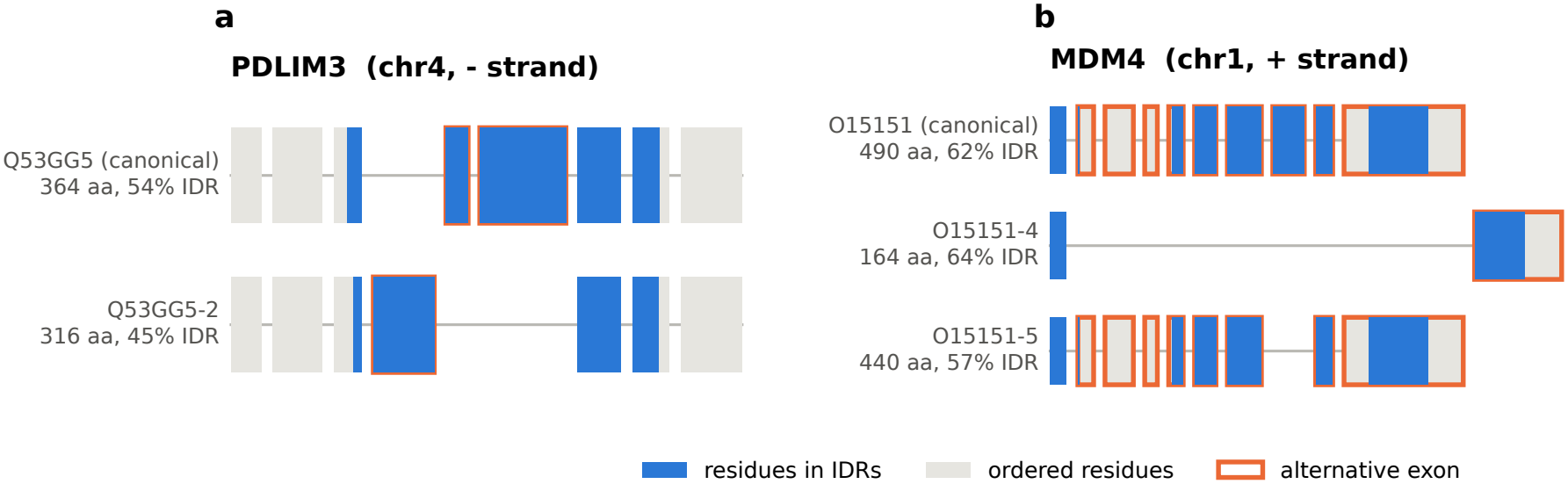
Examples of IDR reorganisation by splicing. Coding exons of each mapped isoform in transcript order, with introns compressed and exon width proportional to the number of encoded residues. Blue, residues in predicted IDRs (*≥*20 residues); grey, ordered residues; orange outline, coding exon present in some but not all of the gene’s isoforms. **a**, PDLIM3: two disordered central exons in the canonical isoform are replaced by a single different disordered exon, rewriting the linker between the PDZ and LIM regions. **b**, MDM4: skipping of a fully disordered internal exon (O15151-5, an internal deletion) and replacement of all downstream exons by an alternative terminal exon (O15151-4, a truncation).

**Extended Data Fig. 6.**
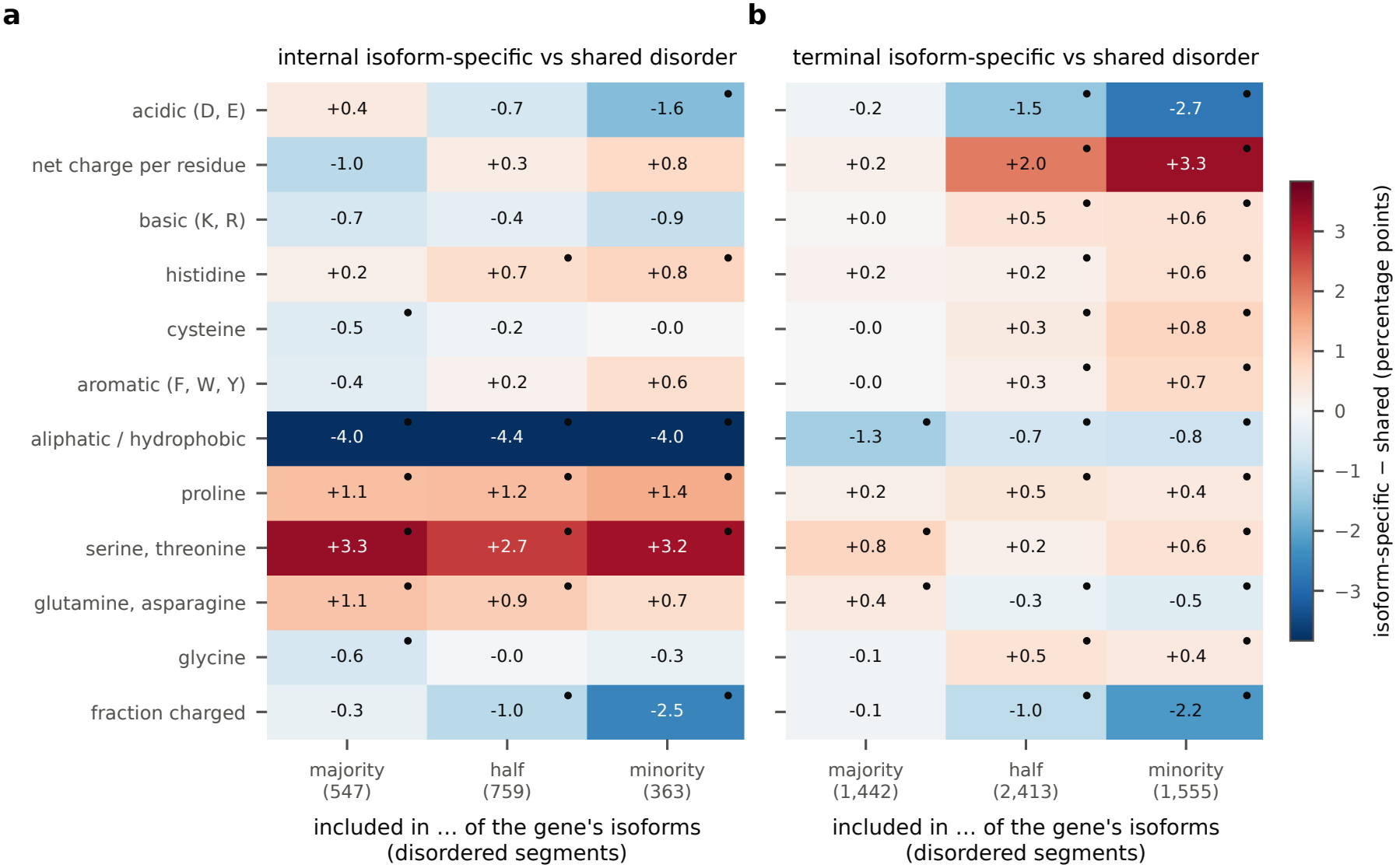
Composition of isoform-specific disorder by inclusion and position. Composition of disordered positions present in only some of a gene’s isoforms compared with the gene’s shared disorder, by inclusion (a majority, half or a minority of isoforms; all isoform-specific positions together are in Fig. 5d). Colour and numbers, difference in percentage points (isoform-specific minus shared, gene-paired); dots, Benjamini–Hochberg *q <* 0.05; number of disordered segments in parentheses. Net charge per residue and fraction charged are multiplied by 100. **a**, Internal isoform-specific disorder. **b**, Terminal isoform-specific disorder.

**Extended Data Fig. 7.**
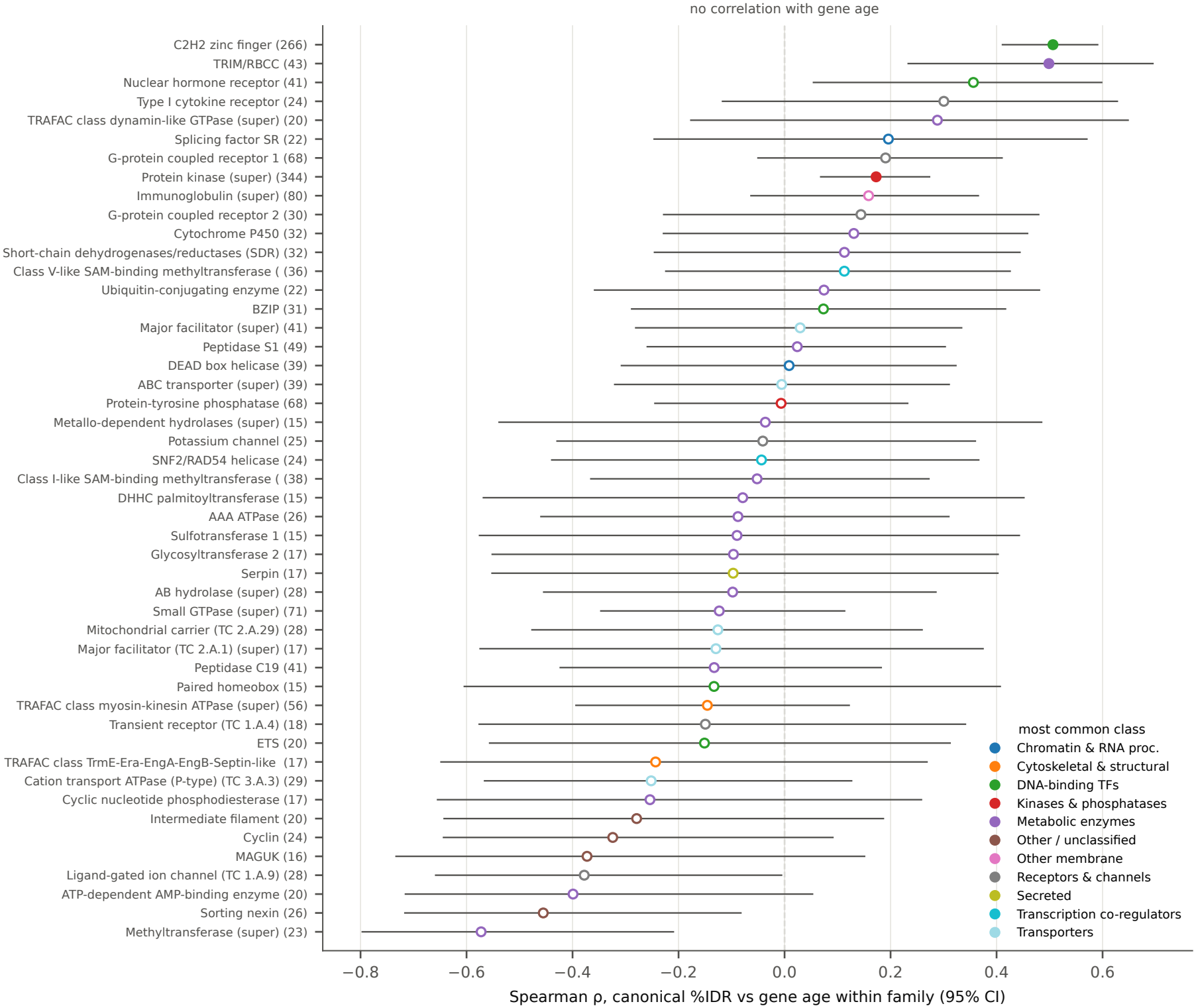
Gene age and disorder within protein families. Spearman correlation of canonical disorder with gene age within each UniProt family with at least 15 dated genes in at least two age bins (95% intervals, Fisher *z*); filled, Benjamini–Hochberg *q <* 0.05; colour, the family’s most common protein class; number of genes in parentheses.

**Extended Data Fig. 8.**
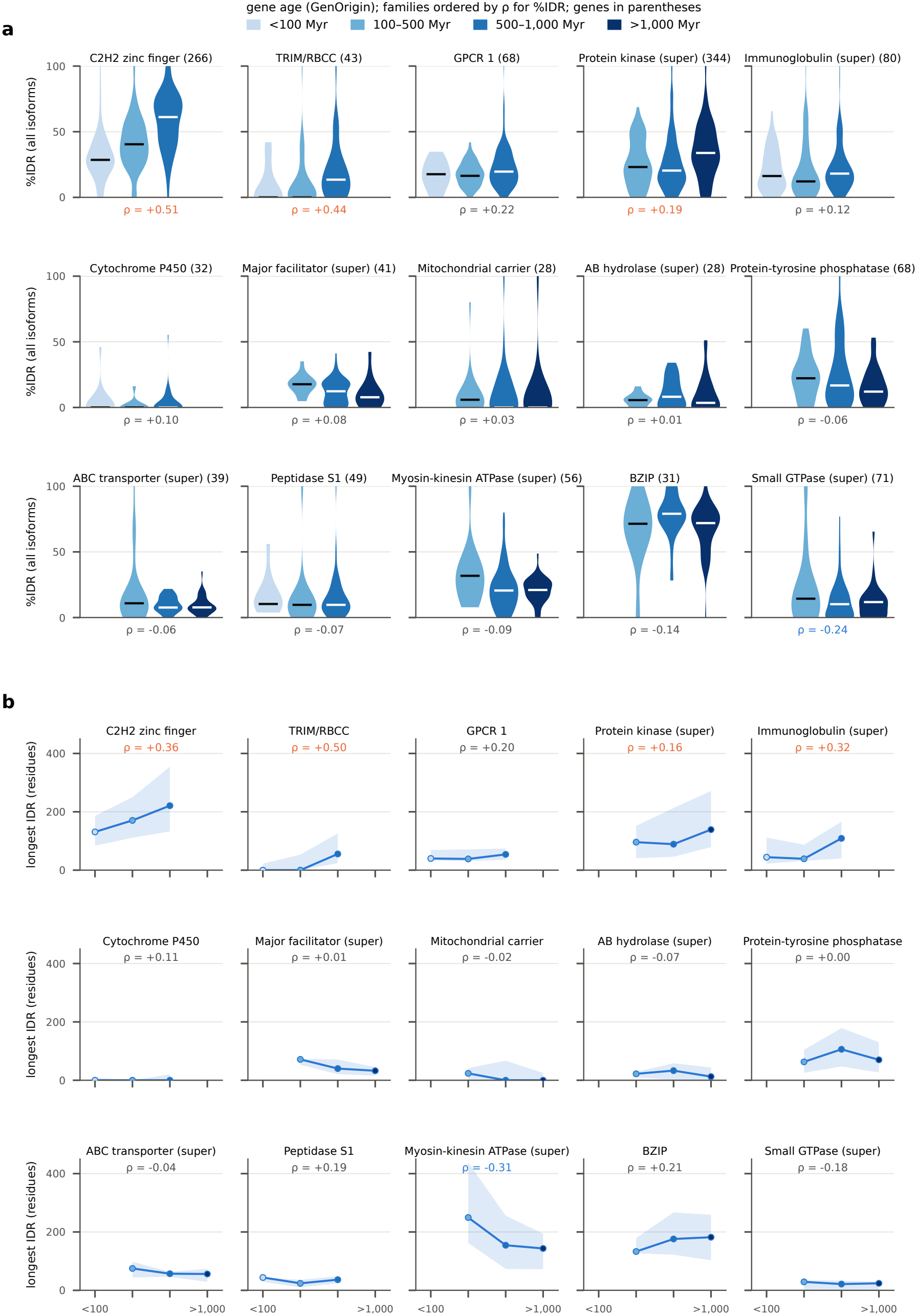
Disorder of all isoforms through gene age within protein families. The 15 UniProt families with at least 15 dated genes and at least five genes in each of three or more age bins, ordered by the correlation for disorder; number of genes in parentheses. Age bins with fewer than five genes are left empty. *ρ*, Spearman correlation of gene age with each gene’s mean over its isoforms (blue, negative; orange, positive; grey, *P ≥* 0.05). **a**, Percentage of residues in IDRs for every isoform (violins over isoforms; line, median). **b**, Length of the longest IDR of every isoform (median, line; interquartile range, band).

**Extended Data Fig. 9.**
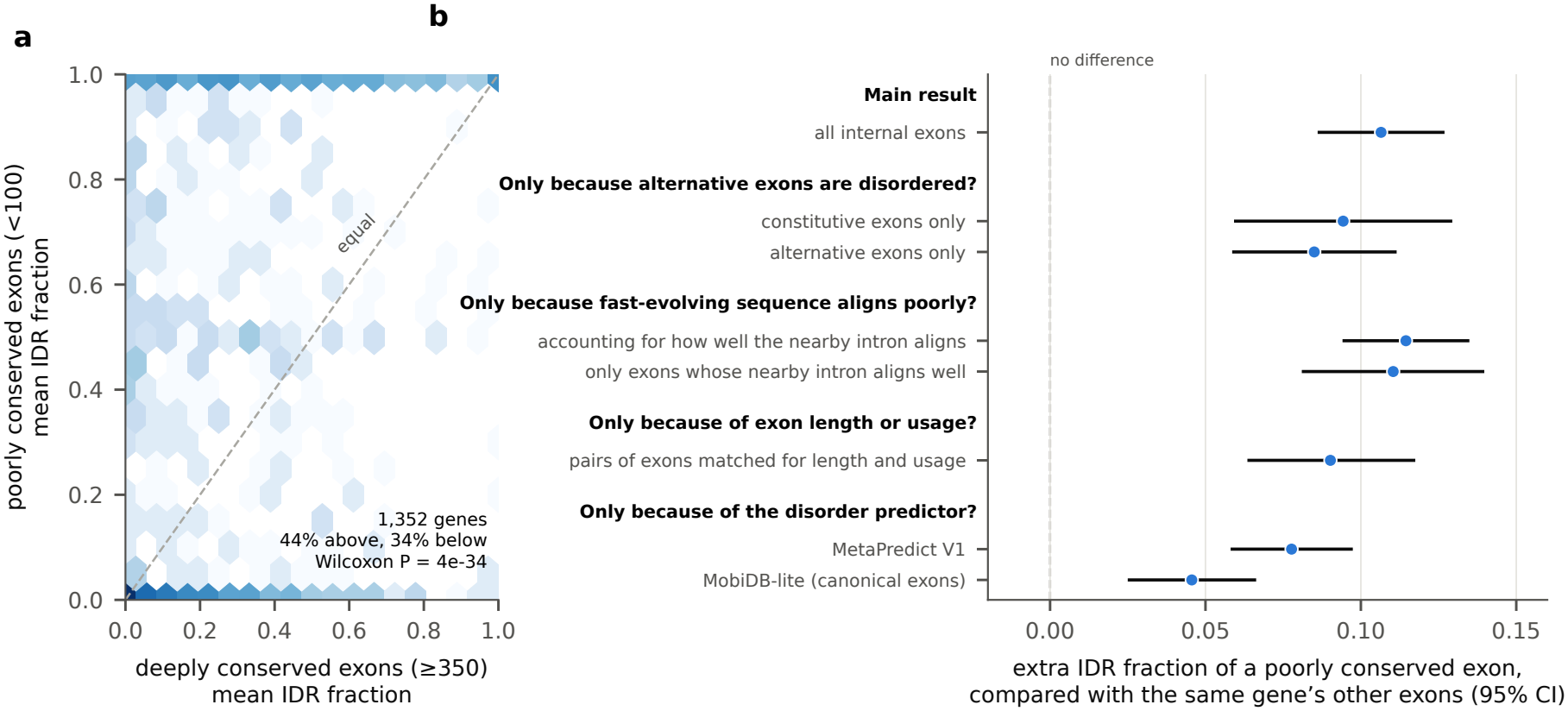
Splice-site conservation: within-gene comparisons. **a**, Genes with both poorly conserved (*<*100 genomes) and deeply conserved (*≥*350) internal exons: mean IDR fraction of each set; dashed line, equality; shading, log gene count. **b**, The within-gene result asked as a series of simple questions. Each point is the extra IDR fraction of exons with a poorly conserved splice site (*<*100 genomes) compared with the other internal exons of the same gene (95% intervals); a value of 0.1 means about 10 more residues in IDRs per 100 residues. Main result: all internal exons. Is it only because alternative exons are disordered? Constitutive or alternative exons alone. Is it only because fast-evolving sequence aligns poorly across species, which could make splice sites look lost? Accounting for how well the 30 intronic bases next to each splice site align, or keeping only exons whose nearby intronic bases align in at least 200 genomes. Is it exon length or usage? Pairs of exons from the same gene matched for length and usage. Is it the disorder predictor? MetaPredict V1 and MobiDB-lite (canonical exons). In every case the difference stays above zero. Adjusting instead for conservation of the exon’s own flanking bases is not used, because that conservation itself falls with disorder (Extended Data Fig. 2a).

**Extended Data Fig. 10.**
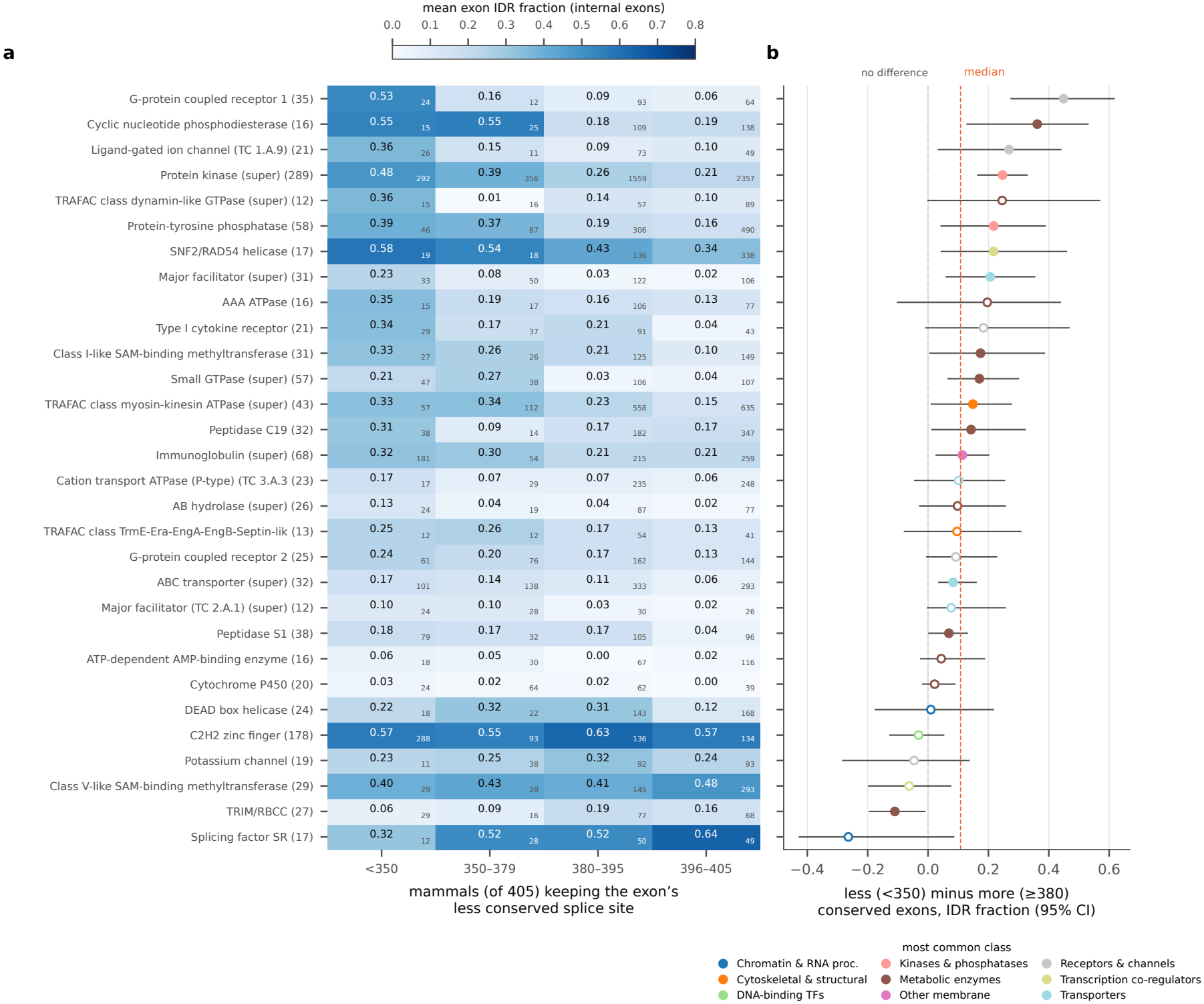
Exon IDR fraction by splice-site conservation within protein families. UniProt families with at least 15 genes and at least ten internal exons in every conservation bin, ordered by the within-family difference; number of genes in parentheses. **a**, Mean IDR fraction of internal exons by the number of 405 mammalian genomes keeping the GT/AG of the exon’s less conserved splice site[21]; small numbers, exons per cell. **b**, Mean IDR fraction of exons conserved in fewer than 350 genomes minus exons conserved in at least 380 genomes of the same family (95% intervals from 500 bootstrap resamples of genes); filled, interval excludes zero; colour, the family’s most common protein class; dashed line, median across families.

